# Non-canonical Wnt signaling promotes directed migration of intestinal stem cells to sites of injury

**DOI:** 10.1101/2021.09.15.460565

**Authors:** Daniel Jun-Kit Hu, Jina Yun, Justin Elstrott, Heinrich Jasper

## Abstract

Tissue regeneration after injury requires coordinated regulation of stem cell activation, division, and daughter cell differentiation, processes that are increasingly well understood in many regenerating tissues. How accurate stem cell positioning and localized integration of new cells into the damaged epithelium are achieved, however, remains unclear. Here we show that enteroendocrine cells coordinate stem cell migration towards a wound in the *Drosophila* intestinal epithelium. In response to injury, EEs release the N-terminal domain of the PTK7 orthologue, Otk, which activates non-canonical Wnt signaling in ISCs, promoting actin-based protrusion formation and ISC migration towards a wound. We find that this migratory behavior is closely linked to ISC proliferation, and that it is required for efficient tissue repair during injury. Our findings highlight the role of non-canonical Wnt signaling in regeneration of the intestinal epithelium, and identify EE-released ligands as critical coordinators of ISC migration.

## Introduction

Migration of somatic stem cells (SCs) has been described as an important phenomenon during regeneration of various tissues^1-3^. Skin SCs, for example, undergo directed migration from the hair follicle to the damaged epidermis during wound healing^4-6^. Hematopoietic and mesenchymal SCs, in turn, are even capable of long-range movement, migrating from the bone marrow to inflamed tissues as part of the immune response^7-10^. The underlying mechanisms regulating and coordinating SC migration, however, have only recently been explored, and technical challenges in imaging living tissue have limited the ability to observe and study this process.

The adult *Drosophila* intestine serves as a powerful model to study intestinal stem cell (ISC) activity and function while, critically, enabling a live imaging platform to directly observe SC behavior in a barrier epithelium^11-16^. ISCs line the pseudo-stratified epithelium and give rise to all the other cell types of the intestine: enteroblasts (EBs, post-mitotic precursor cells), enterocytes (ECs, differentiated cells with scaffolding and nutrient absorption roles), and enteroendocrine cells (EEs, differentiated cells with secretory roles)^13,17^. ISCs are largely quiescent during homeostasis, but are activated to divide in response to tissue damage or during tissue growth^11,18,19^. Studies of ISC dynamics during regeneration have largely been constrained to static analysis of fixed tissue, but recent innovations in imaging live, wholemount intestinal explants has expanded the ability to investigate these processes in real time, providing insights into symmetric and asymmetric ISC divisions, intracellular calcium signaling, cell loss, and cell fate determination and differentiation in the ISC lineage^11-16^.

Cell migration is an actin-based process in which members of the Rho family of GTPases establish polarity at the leading edge by activating the Arp2/3 complex and mDia^20-23^. These proteins, in turn, polymerize actin to form lamellipodia and filopodia, directing forward motion through forces generated from actin flow and actomyosin contractility^24-26^. During development and morphogenesis, non-canonical Wnt signaling links extracellular cues to actin rearrangement through the interaction of Wnt ligands with the cell surface receptors Frizzled (Fz), Ptk7 (Otk in *Drosophila*), and Ror2 (Ror in *Drosophila*)^27-33^. This interaction activates intracellular Dishevelled (Dsh in *Drosophila*, Dvl in mammals), which leads to downstream activation of Rho GTPases and actin rearrangement^34-36^. The extent in which this pathway plays a role in adult SC migration and regeneration remains unresolved.

Here, we demonstrate that *Drosophila* ISCs rapidly initiate migration after enteropathogen infection and after localized tissue damage by laser ablation. This process is mediated by a signaling cascade relying on matrix-metalloproteinase (MMP) induction and Otk expression in EEs at the wound site, which in turn activates non-canonical Wnt signaling in ISCs, promoting actin-dependent formation of lamellipodia, and migration of ISCs to the wound area. Impairing ISC migration hinders ISC proliferation as well as effective intestinal regeneration following tissue damage, and sensitizes animals to death by enteropathogen infection. We propose that MMP-mediated cleavage of Otk in EEs at the wound is a critical signal promoting ISC migration toward the site of epithelial injury, ensuring efficient regeneration.

## Results

### Intestinal stem cells exhibit migratory behavior after tissue damage

To visualize ISC behavior in response to damage, we imaged live, wholemount fly intestines *ex vivo* (Fig. 1a), a system we have previously utilized to directly observe ISC mitoses, EB differentiation, and intracellular calcium dynamics^11,12^. ISCs were identified by specific and inducible expression of cytoplasmic eYFP using the ISC/EB driver *escargot::Gal4* in combination with EB-specific expression of the Gal4 inhibitor Gal80 (Su(H)::Gal80), and ubiquitous expression of temperature-sensitive Gal80 (tub::Gal80^ts^)^37,38^. Flies were infected with *Erwinia carotovora carotovora 15* (*Ecc15*), a gram-negative bacterium that damages differentiated enterocytes, to induce a regenerative response (Fig. 1a)^39,40^. Under homeostatic conditions, ISCs were largely immotile, with little change in morphology (Fig. 1b, Supplementary Fig. 1a, and Supplementary Video 1). In stark contrast, after 16 hours post-*Ecc15* infection, ISCs were far more dynamic (Fig. 1b, Supplementary Fig. 1a, and Supplementary Video 2). Over the 2.5hr timelapse, ∼80% of ISCs formed protrusions resembling a leading edge, indicative of migratory behavior. 57% of the ISCs that formed these protrusions fully migrated, as determined by translocation of the cell body of at least 3µm. The migratory capability of ISCs in mock-treated or infected intestines was also compared by tracking the location of the cell body and the tip of the protrusion, or the most distal region of the cell cortex if a protrusion did not form (Fig. 1b and Supplementary Fig. 1a). When the x,y coordinates of protrusions or cell bodies at each time point were plotted on a Cartesian plane, both the cell body and the most distal part of the cell cortex travelled dramatically farther in ISCs from infected intestines than in controls. We asked whether other cell types were also capable of migration, and used the Su(H) and prospero promoters to identify post-mitotic precursor EBs and differentiated EEs respectively. Only minimal protrusions and movement were observed in these cell types after *Ecc15* infection (Supplementary Fig. 1b and Supplemental Video 3-4) suggesting that, in contrast to ISCs, more differentiated cells are integrated statically into the epithelium.

**Figure 1:**
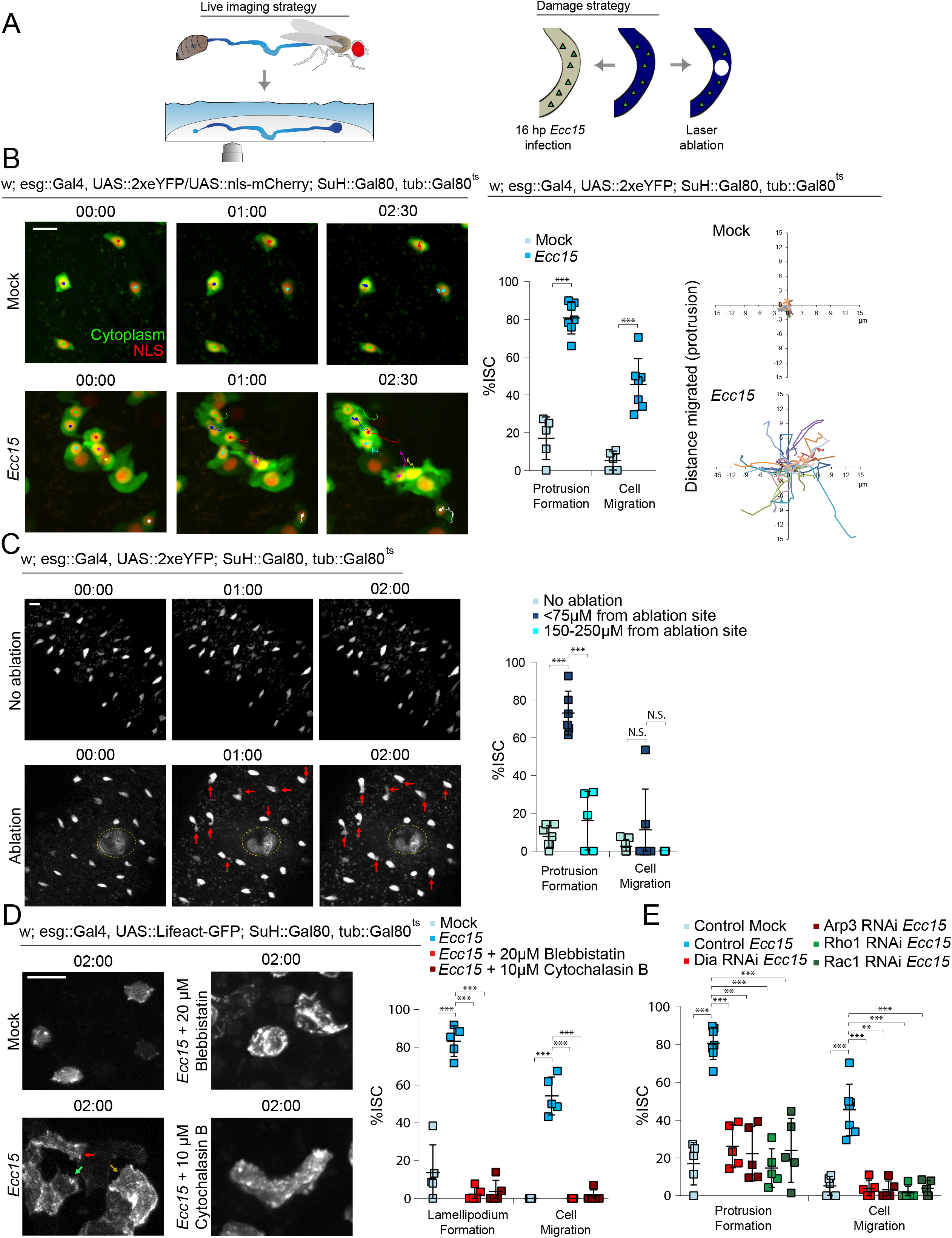
ISCs undergo migration following tissue damage. A. Diagram representing *ex vivo* live imaging and damage strategy of *Drosophila* intestine. B. Montage and quantification of protrusion formation and migration of ISCs in mock-treated and *Ecc15*-infected intestines. Montage tracks measured from the center of the nucleus, with starting x,y coordinate of each ISC normalized to 0,0. C. Montage and quantification of protrusion formation and migration of ISCs in undamaged intestine, near the wound (≤75µm) of laser-ablated intestines, and far away from the wound (150-250µm) of laser-ablated intestines. Red arrows indicate protrusions and dotted line indicates ablation site. D. ISCs formed lamellipodia after *Ecc15*-infection. Lamellipodia formation and migration of ISCs dramatically decreased after inhibition of actomyosin function with small molecule inhibitors. Arrows indicate lamellipodia. E. Expression of RNAi for proteins involved in regulating actin assembly decreased protrusion formation and migration of ISCs. Mock-treated and *Ecc15*-infected controls were taken from Figure 1B, as they served as genetic controls and were performed in parallel. mean ± SD; n≥5 flies; N.S. = not significant, ***P<0.001, based on Student’s t-test (B) and one-way ANOVA with Tukey test (C,D,E). Scale bar = 10µm. Timestamp indicated as hours:minutes. See also Supplemental Figure 1, 2, and 3, Supplemental Video 1-11.

*Ecc15* causes wide-spread damage in the intestinal epithelium, and this paradigm was thus not suited for an examination of directed migration of ISCs. To generate a more localized epithelial wound instead, we utilized high-powered lasers with 2-photon microscopy. In addition to creating a localized ∼30µm wound (Fig. 1c), this strategy also enables observation of ISC behavior immediately after damage. 73% of ISCs within 75µm of the wound formed protrusions, compared to 8% in undamaged tissue (Fig. 1c, Supplemental Fig. 1c, and Supplemental Video 5-6). This response was dependent on proximity to the wound as the majority of ISCs located farther away from the wound (150-250µm) did not form protrusions (Fig. 1c).

Despite forming protrusions, the cell bodies of most ISCs within 75µm of the wound did not translocate in the first 2.5 hours post ablation, potentially because the migratory response continues beyond this timeframe (the structural integrity of the ablated tissue began to deteriorate after three hours of live imaging, making longer imaging durations challenging). Nonetheless, we observed that the protrusions of ∼81% of ISCs formed towards the wound, suggesting that migratory behavior was polarized towards the site of injury (Supplemental Fig. 2a). To assay for directionality of migration at later time points while avoiding potential toxicity from frequent 2-photon microscopy, we imaged ablated guts immediately after ablation and 4.5 hours later. We observed that most ISCs were located closer to the wound after 4.5 hours whereas ISCs from unablated tissue rarely changed their position (Supplemental Fig. 2b). Furthermore, when damaged guts were cultured for 4.5 hours before fixation, we observed that ISCs had accumulated around the periphery of the wound, and that there was a greater number of ISCs in a 40µm radius around the ablation site than within the same area of the contralateral side of the intestine (Supplemental Fig. 2c; there was approximately the same number of ISCs on contralateral sides of undamaged intestines). Importantly, we did not observe any mitoses within the first 2.5 hours after ablation (duration of live experiments) or at 4.5 hours post ablation (fixed experiments), suggesting that the accumulation of ISCs around the wound was a result of directed migration rather than generation of additional ISCs by mitosis.

### ISC migration is dependent on the formation of actin-based lamellipodia

The presence of protrusions in migratory ISCs suggests that migration is driven by lamellipodia formation. To directly visualize whether a lamellipodium did indeed form, GFP-labelled LifeAct^14^ was expressed in ISCs. Under homeostatic conditions, cortical actin was observed around the periphery of ISCs (Fig. 1d and Supplemental Video 7). After *Ecc15* infection, actin becomes enriched at the leading edge, and the formation of fliopodia and the lamellipodium were observed (Fig 1d, Supplemental Fig. 3a, and Supplemental Video 8). Following extension of the lamellipodium, translocation of the cell body and retraction of the trailing edge then occurred. There were zero observed cases of translocation of the cell body without lamellipodium formation. To confirm that lamellipodia formation was actin-dependent, we utilized small molecule inhibitors to block actin polymerization or actomyosin contraction. Addition of either Cytochalasin B or Blebbistatin (small molecule inhibitors for actin network assembly and myosin II function respectively)^41,42^ abolished lamellipodia dynamics and ISC migration (Fig. 1e and Supplemental Video 9). To further test a role of the lamellipodium in ISC migration, we examined proteins involved in regulating actin dynamics, and thus in lamellipodium formation. We tested the role of Dia (mDia1 in mammals) and the Arp2/3 complex, which promote actin nucleation and branching, as well as their upstream regulators, the GTPases Rho1 (RhoA in mammals) and Rac1^20,43^. Knocking down Dia, Arp3, Rho1, or Rac1 in ISCs of *Ecc15*-infected flies resulted in a significant decrease in protrusion formation and ISC migration compared to control flies (Fig. 1e, Supplemental Fig. 3b, and Supplemental Video 10).

To better understand the mechanism(s) promoting translocation of the cell body, specifically the nucleus, we investigated the roles of nesprins in facilitating nuclear movement. Nesprins are part of the LINC complex that tether the cytoskeleton to the nuclear envelope, enabling nuclear positioning^44^. We depleted Klarsicht (Klar), a *Drosophila* nesprin, in ISCs from infected intestines, and found that the percentage of cell bodies that remained stationary increased compared to control flies (Supplemental Fig. 3c and 4b, and Supplemental Video 11). Despite an inability to translocate the cell body, many of the ISCs after Klar depletion were still capable of forming protrusions, exhibiting a more elongated morphology as the leading edge often continued to extend. A Klar-mediated connection of the nucleus to the cytoskeleton is thus a critical component of the machinery executing ISC migration.

Altogether, these data support the idea that, in response to tissue damage, ISCs exhibit lamellipodia-dependent migration. Because ISCs are largely immotile under homeostatic conditions, ISC migration is likely part of the regenerative response to damage. To test this hypothesis, we sought to identify signaling mechanisms that may mediate the damage-induced activation of lamellipodia formation. Due to its role in cytoskeletal rearrangements and cell migration during development, we focused on the non-canonical Wnt signaling pathway^34,36^.

### Non-canonical Wnt signaling promotes ISC migration

To test whether non-canonical Wnt signaling is required for ISC migration, we knocked down components of the pathway in ISCs, before infecting flies with *Ecc15*. Infection-induced cellular protrusions, as well as ISC migration were no longer observed after depleting Dsh, Otk, or the second *Drosophila* Fz, Fz2, while knock down of the second Drosophila Ptk7 orthologue, Otk2, or Fz had no effect (Fig. 2a, Supplemental Fig. 4a-4b, and Supplemental Video 12-13). Non-canonical Wnt signaling is also required for protrusion formation after laser injury, as knockdown of Dsh or Otk reduced protrusion formation (Fig. 2b and Supplemental Video 14-15). Depleting Dsh or Otk in ablated tissue resulted in a loss of ISC accumulation around the wound at 4.5 hours post ablation, further supporting a role for non-canonical Wnt signaling during directed migration (Supplemental Fig. 4c). Protrusion formation was also reduced after treatment with the porcupine inhibitor LGK974^45^, which inhibits release of Wnt ligands, indicating that secreted Wnt ligands play a critical role in inducing or maintaining migration of ISCs (Fig. 2c and Supplemental Video 16). Wingless (Wg), a *Drosophila* Wnt, has been previously reported to be secreted from enteroblasts following damage of the *Drosophila* intestine, acting as an important regulation of ISC proliferation during tissue regeneration^46^. We tested whether EBs could also serve as a source of Wg to promote ISC migration after tissue damage and observed that, indeed, depleting Wg in EBs reduced the percentage of ISCs exhibiting migratory behavior (Supplemental Fig. 4d).

**Figure 2:**
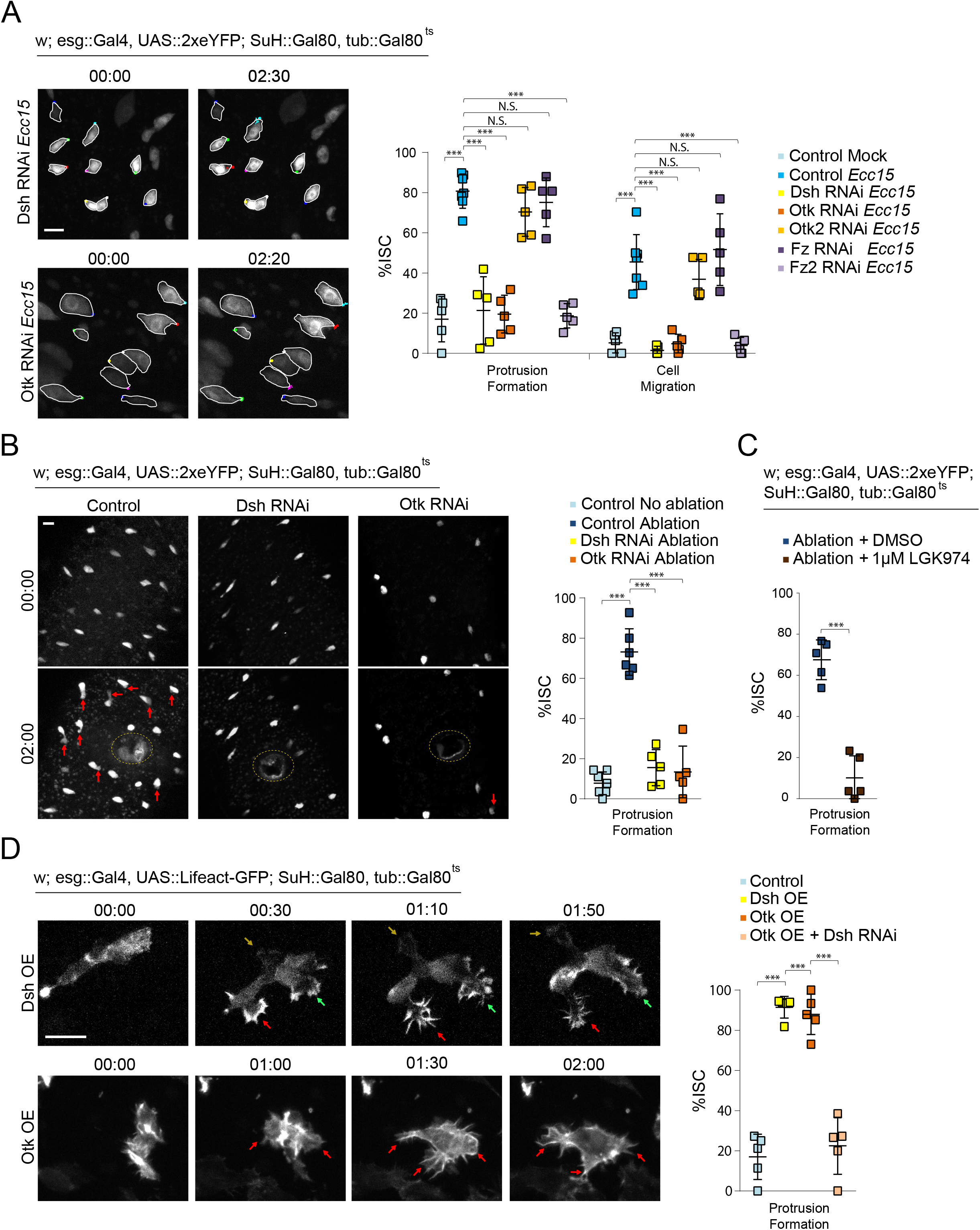
Non-canonical Wnt activity promotes ISC migration. A. Quantification of ISC migratory behavior after genetic perturbation, and montage after depleting Dsh and Otk. ISCs outlined in white, and montage tracks measured from the most distal region of the cell cortex. Mock-treated and *Ecc15*-infected controls were taken from Figure 1B, as they served as genetic controls and were performed in parallel. B. Montage and quantification of ISC protrusion formation after laser ablation in Dsh- and Otk-depleted intestines. Red arrows indicate protrusions and dotted line indicates ablation site. Undamaged and ablated controls were taken from Figure 1c, as they served as genetic controls. C. Quantification of ISC protrusion formation in ablated intestines after inhibiting Wnt secretion versus DMSO-treated controls. D. Montage and quantification of lamellipodia formation after overexpressing Dsh or Otk. Lamellipodia formation induced by Otk overexpression is suppressed after depleting Dsh. Arrows indicate protrusions. mean ± SD; n≥5 flies; N.S. = not significant, ***P<0.001, based on one-way ANOVA with Tukey test (A,B,D) and Student’s t-test (C). Scale bar = 10µm. Timestamp indicated as hours:minutes. See also Supplemental Figure 4, Supplemental Video 12-17.

Activation of non-canonical Wnt signaling is also sufficient for the induction of migratory behavior in ISCs, as overexpressing either myc-tagged Dsh or GFP-tagged Otk in ISCs in unchallenged, undamaged guts caused dramatic rearrangement of the actin cytoskeleton (Fig. 2d and Supplemental Video 17). The Otk-GFP signal is too weak to observe without an anti-GFP antibody, and is undetectable compared to the brighter GFP-tagged LifeAct. ISCs continually formed filopodia, often with multiple, seemingly non-polarized protrusions, which extended from different regions around the cell cortex. We confirmed that Dsh acts downstream of Otk in this context, as knockdown of Dsh while overexpressing Otk in ISCs impaired protrusion formation (Fig. 3d). Importantly, canonical Wnt signaling appears not to be involved in ISC migration, as knockdown of the canonical Wnt pathway transducers Armadillo (Arm; ß-catenin in mammals) or Pangolin (Pan; TCF in mammals)^47,48^ did not affect ISC migration after *Ecc15* infection (Supplemental Fig. 4e).

**Figure 3:**
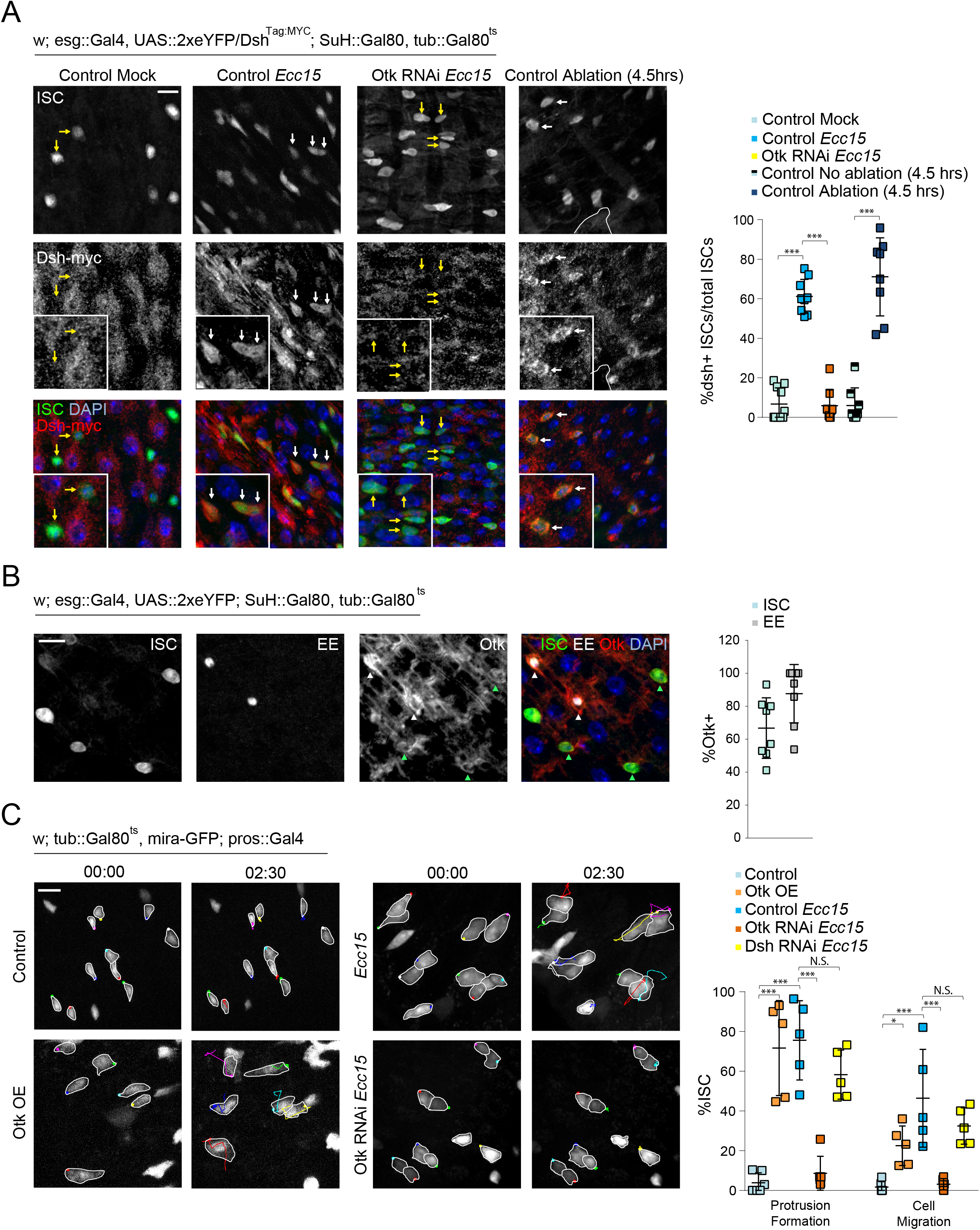
Localization of non-canonical Wnt pathway proteins, and role of EE-derived Otk in ISC migration. A. Staining for endogenously myc-tagged Dsh reveals cortical decoration in ISCs after tissue damage, but absent or sparse localization in ISCs from undamaged tissue. Cortical Dsh is lost after Otk depletion in ISCs of damaged tissue. White outline indicates ablation site. White arrows indicate Dsh+ ISCs and yellow arrows indicate Dsh-ISCs. B. Staining for Otk reveals expression predominantly in ISCs (green arrowhead) and EEs (white arrowhead). C. Montage and quantification of ISC migratory behavior after modulating Otk and Dsh expression in EEs. ISCs outlined in white, and montage tracks measured from the leading edge or, in cells that did not form protrusions, the most distal region of the cell cortex. mean ± SD; n≥8 flies (A,B), n≥5 flies (C); N.S. = not significant, *P<0.05, ***P<0.001, based on one-way ANOVA with Tukey test. Scale bar = 10µm. Timestamp indicated as hours:minutes. See also Supplemental Figure 5, Supplemental Video 18-19.

To further confirm and explore the role of non-canonical Wnt signaling in promoting ISC migration after damage, we assessed the protein expression and localization of Dsh and Otk. Dsh localization to the cell cortex is a hallmark of non-canonical Wnt signaling activity^49-51^. Using a fly line expressing myc-tagged Dsh under control of its endogenous promoter^52^, we found that Dsh was undetectable in ISCs from mock-treated, undamaged intestines, potentially either due to low expression or sparse localization at the cell membrane (Fig. 3a). Strikingly, after damage from either 16 hours post *Ecc15* infection, or 1 or 4.5 hours post laser ablation, Dsh became strongly localized to the cell membrane of ISCs, indicating an activation of non-canonical Wnt signaling after damage (Fig. 3a and Supplemental Fig. 5a). Consistent with a role of Otk in this activation, Otk depletion in damaged intestines was sufficient to abolish Dsh localization to the ISC membrane.

To examine whether Otk could be detected at the cell membrane of ISCs, we generated antibodies recognizing the intracellular, C-terminal region of Otk, which successfully detected overexpressed GFP-tagged Otk at the cell membrane (Supplemental Fig. 5b). Otk was not detected in EB or EC cells, but was present in both ISCs and EEs (Fig. 3b). EEs were previously reported to play important roles in facilitating ISC activity^53^, and we tested whether EEs could also regulate ISC migration through Otk. Overexpression of Otk in EEs using the EE-specific driver prospero::Gal4, promoted protrusion formation in ISCs (detected in this experiment using miranda::GFP^54^), suggesting a non-autonomous role of Otk (Fig. 3c and Supplemental Video 18). This effect seemed to be EE specific as overexpression of Otk in the EBs or ECs did not affect ISC migratory rates (Supplemental Fig. 5c). Conversely, depleting Otk in EE cells of *Ecc15*-infected guts was sufficient to decrease protrusion formation in ISCs, and impair ISC migration (Fig. 3c and Supplemental Video 19). However, the role of EE-derived Otk in ISC migration did not seem to involve non-canonical Wnt signaling within EEs themselves as EE-specific depletion of Dsh did not affect ISC migration (Fig. 3c and Supplemental Fig. 5d). Because the Otk antibody also seemed to stain the muscle layer (Fig. 3b), we tested whether depleting Otk from the muscle layer had any effect on ISC migration, and found no impairment of migratory behavior (Supplemental Fig. 5e).

### EE-derived shedding of Otk is required for ISC migration

Previous studies in mammals have found that the extracellular, N-terminal region of Ptk7 can be cleaved from the cell membrane by MMP and ADAM metalloproteinases to promote migration in cancer cells^55-57^. We examined whether this ‘shed’ form of Ptk7 was sufficient to induce migratory behavior in ISCs. Undamaged intestines were incubated with a recombinant, extracellular fragment of mouse Ptk7 (Met1-Glu683). Within an hour post incubation, the majority of ISCs underwent rapid filopodia formation with highly dynamic protrusions, phenocopying Otk overexpression (Fig. 4a and Supplemental Video 20). The effect of extracellular Ptk7 was also dependent on non-canonical Wnt signaling in ISCs, as depleting Otk or Dsh in ISCs reduces protrusion formation (Fig. 4a and Supplemental Video 21).

**Figure 4:**
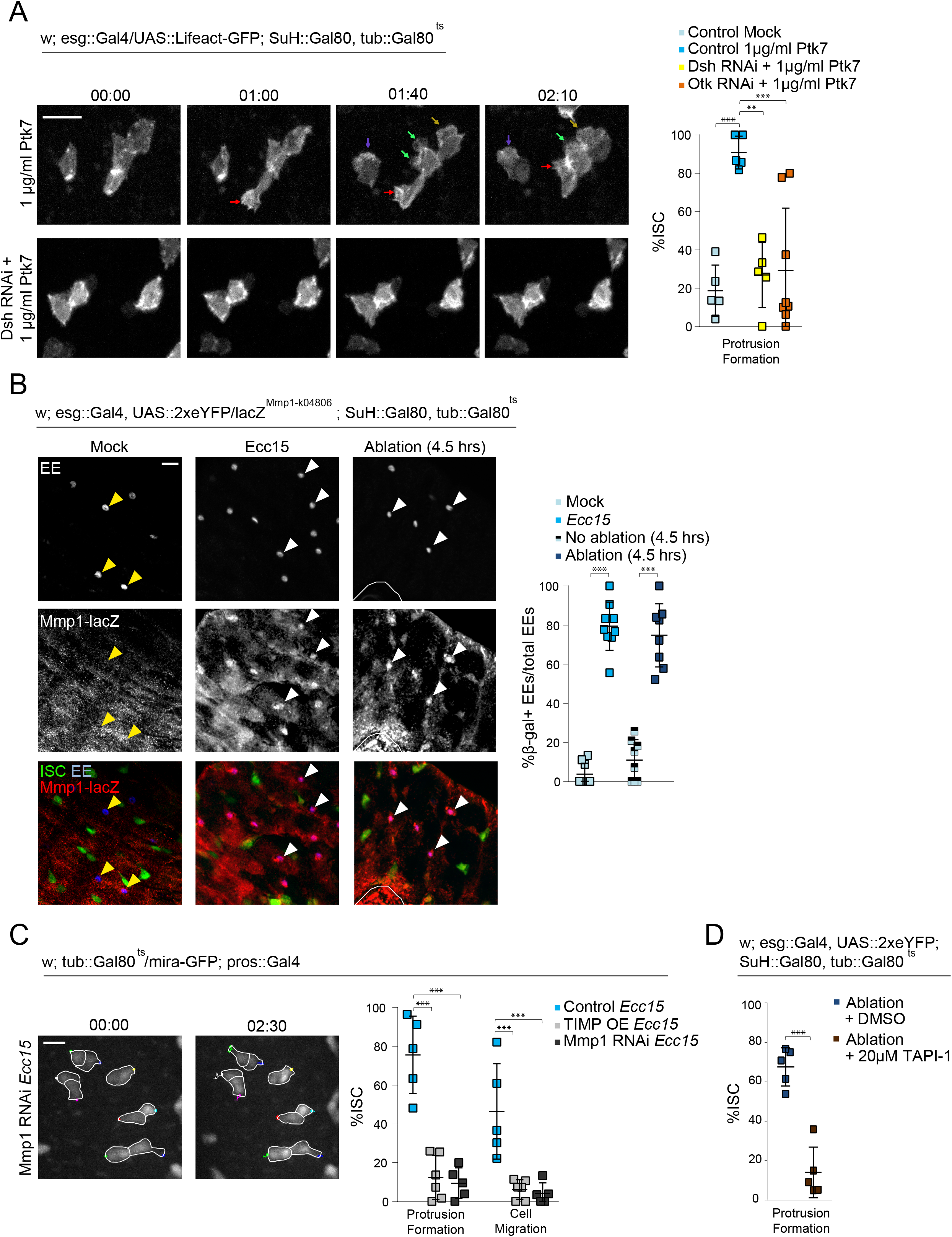
Injury-induced Mmp1 in EEs mediates ISC migration. A. Montage and quantification of actin dynamics in ISCs after Ptk7 treatment in control, Dsh^RNAi^, and Otk^RNAi^ intestines. Arrows indicate protrusions. B. Staining for beta Galactosidase in Mmp1-lacZ reporter lines reveal Mmp1 expression in EEs after tissue damage. White arrowheads indicate β-gal+ EEs and yellow arrowheads indicate β-gal-EEs. White outline indicates ablation site. C. Montage and quantification of ISC migratory behavior after disrupting Mmp1 activity. *Ecc15*-infected controls were taken from Figure 3C, as experiments were performed in parallel. ISCs outlined in white, and montage tracks measured from the most distal region of the cell cortex. D. Quantification of ISC protrusion formation in ablated intestines after inhibiting MMP activity versus DMSO-treated controls. mean ± SD; n≥5 flies (A,C,D), n≥8 flies (B); N.S. = not significant, **P<0.01, ***P<0.001, based on one-way ANOVA with Tukey test (A,C) and Student’s t-test (B,D). Scale bar = 10µm. Timestamp indicated as hours:minutes. See also Supplemental Figure 6, Supplemental Video 20-23.

Because extracellular Ptk7 was sufficient to drive migratory behavior, we hypothesized that, in response to damage, Otk in EEs may be cleaved by MMPs to promote ISC migration. To test this idea, we assessed Mmp1 expression using an Mmp1-lacZ reporter line^58^. While Mmp1 expression was sparse in undamaged tissue, beta Galactosidase expression was detected strongly and specifically in EEs after *Ecc15* infection or laser ablation (Fig. 4b and Supplemental Fig 6a). We were able to recapitulate these results using antibodies against Mmp1, and found that Mmp1 staining was normally absent in EEs in uninjured tissue, but becomes detectable after bacterial infection (Supplemental Fig. 6b). We inhibited Mmp1 activity specifically in EEs by overexpression of its inhibitor, TIMP, or by depletion of Mmp1, and observed a drastic reduction of protrusion formation and migratory ability of ISCs after *Ecc15* infection (Fig. 4c and Supplemental Video 22). Furthermore, treatment of intestines with TAPI-1, an ADAM and MMP inhibitor^59^, prior to laser ablation, decreased protrusion formation in ISCs (Fig. 4d and Supplemental Video 23).

To better visualize cleavage of Otk, we overexpressed in EEs full-length Otk tagged with GFP at the intracellular C-terminus, and immunostained for the extracellular, N-terminal domain of Otk. Both N-terminal and C-terminal domains were present in EEs in uninjured guts, but N-terminal Otk was no longer present after *Ecc15* infection (Fig. 5a). Crucially, C-terminal Otk was still present, suggesting Otk is still trafficked to the EE membrane (Fig. 5a). The N-terminal domain of endogenous Otk was also detectable in EEs in homeostatic conditions (albeit at much lower intensity than the over-expressed molecule), but was absent after *Ecc15* infection (Fig. 5b). This loss of N-terminal Otk was no longer observed after EE-specific depletion of Mmp1 (Fig. 5b). Interestingly, in ISCs, the N-terminal domain of Otk was detectable in both uninjured and infected guts, suggesting that Otk is not cleaved in ISCs (Fig. 5b). Overall, these data provide evidence for a crucial role of Mmp1-dependent shedding of Otk in EE cells to activate ISC migration during the regenerative response.

**Figure 5:**
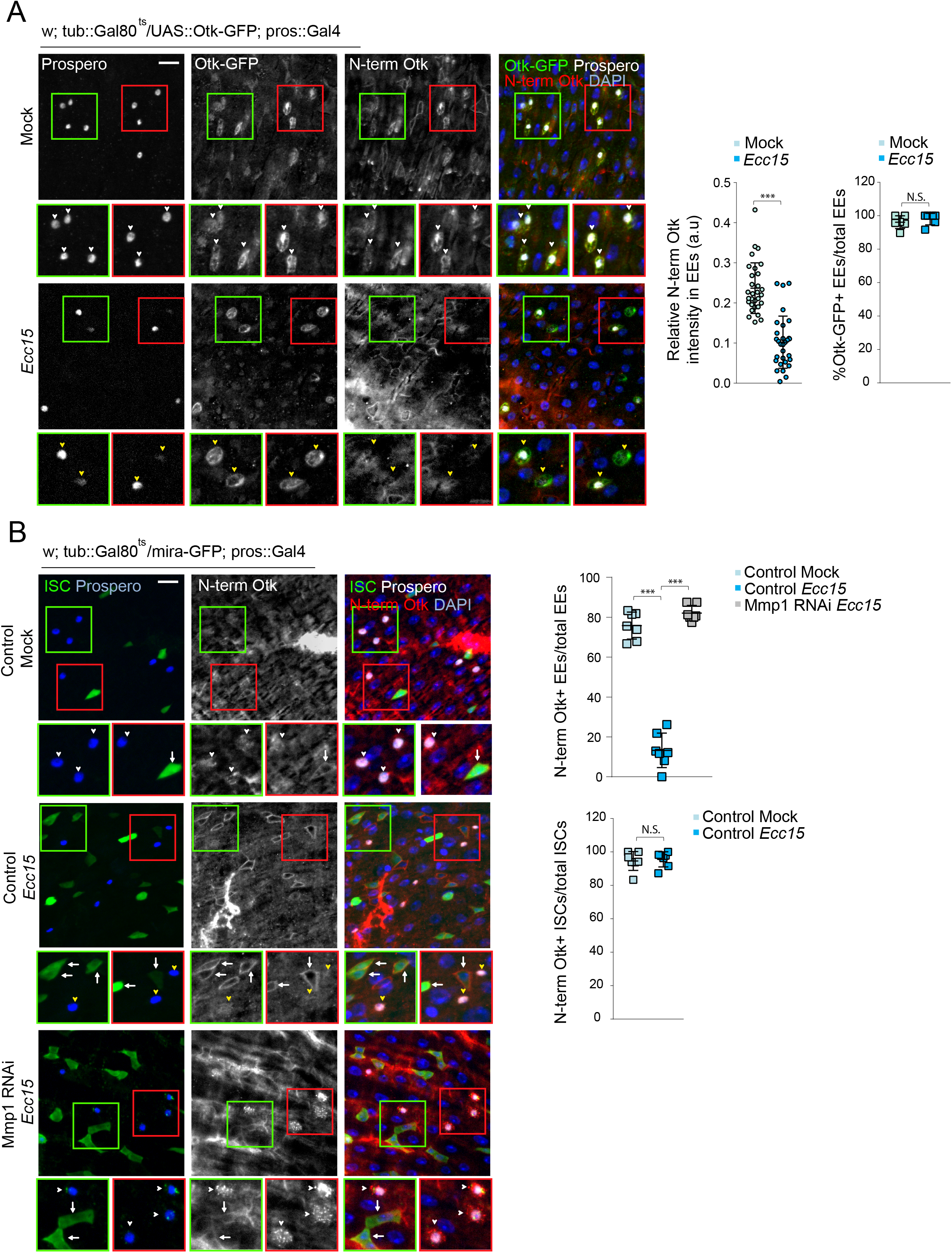
Extracellular Otk is cleaved in EEs by Mmp1 after tissue injury. A. Full-length Otk with a GFP tag on the C-terminal, intracellular region was overexpressed specifically in EEs to visualize intracellular Otk. Antibodies against the N-terminal, extracellular Otk was used to visualize extracellular Otk. Both N-terminal and C-terminal regions of Otk were detected on the EE cortex in undamaged tissue (white arrowhead), but only C-terminal Otk was detected after tissue damage (yellow arrowhead). B. N-terminal Otk was detected in EEs from undamaged tissue (white arrowhead), but was largely absent in EEs from damaged tissue (yellow arrowhead). Depleting Mmp1 in EEs from damaged tissue prevented the loss of N-terminal Otk (white arrowhead). N-terminal Otk was detected in ISCs from both undamaged and damaged tissue (white arrows). mean ± SD; n=30 cells from 6 flies (A), n=6 flies (A), n=7 flies (B); N.S. = not significant, ***P<0.001, based on Student’s t-test (A,B) and one-way ANOVA with Tukey test (B). Scale bar = 10µm.

### Impairing ISC migration decreases regeneration rates

To determine whether migration is indeed part of the regenerative response, we tested whether impairing ISC migration would affect tissue regeneration. We first assessed whether migration of ISCs was coupled to their proliferative activity. Depleting Dsh or Otk decreased the number of mitotic cells after 16 hours of *Ecc15* infection (Fig. 6a). Because the non-canonical Wnt pathway plays multiple signaling roles, we also depleted Klar to more specifically target the mechanics of cell migration. Similar to Dsh and Otk, depleting Klar reduced the number of mitotic figures after infection, suggesting a coupling between cell cycle activation and cell migration (Fig. 6a). Proliferative activity is normally very low under homeostatic conditions, and depleting Dsh, Otk, or Klar did not significantly decrease PH3 numbers in uninjured guts (Supplemental Fig. 7a). As an additional assay to test the effect of reduced ISC migration on cell cycle progression, we used an esg-FlipOut lineage tracing system to generate clones^60^. Indeed, disrupting ISC migration by Dsh, Otk, or Klar RNAi reduced the number of cells per clone after *Ecc15* infection (Supplemental Fig. 7b).

**Figure 6:**
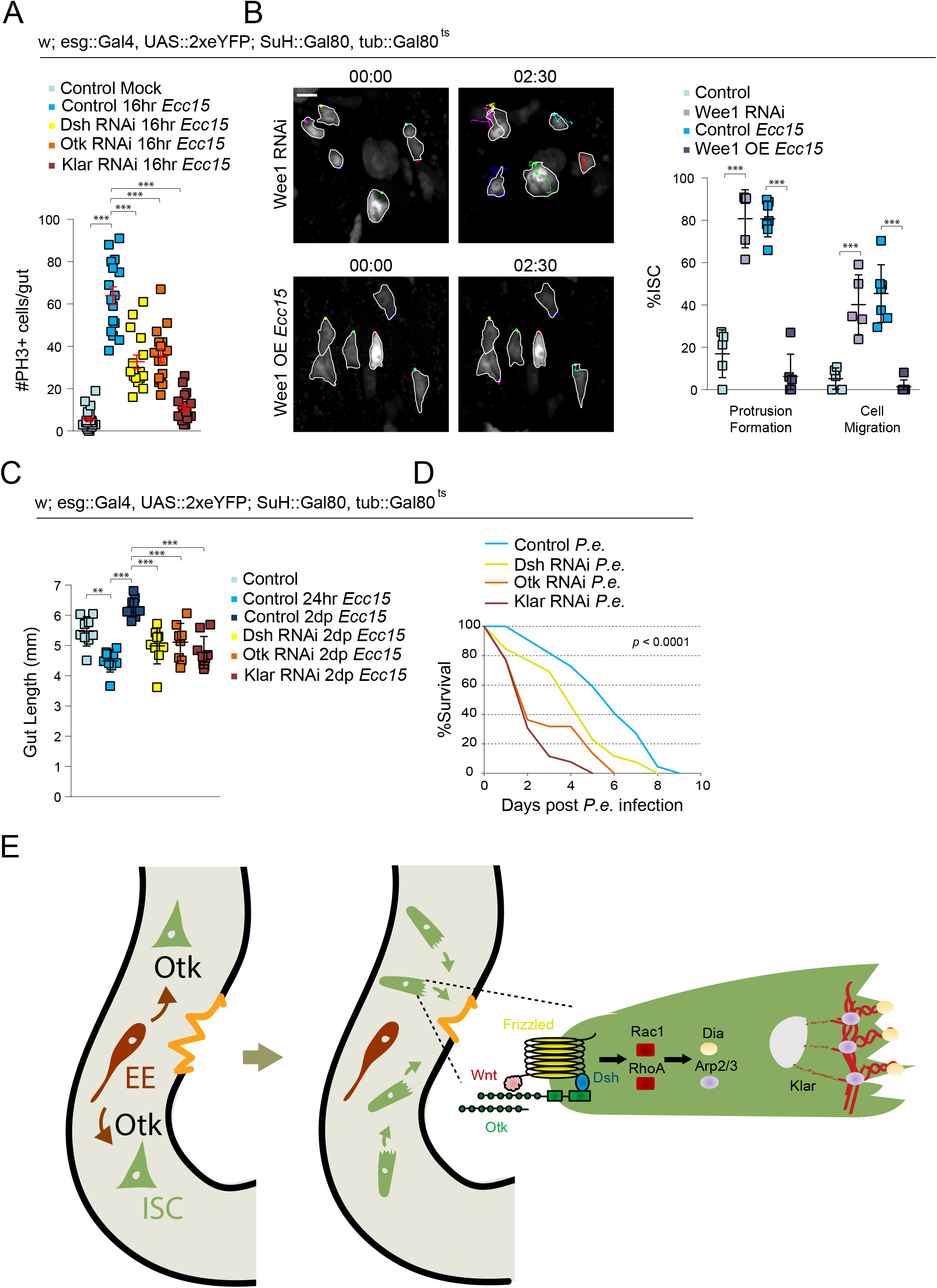
Disrupting ISC migration impairs cell cycle progression and tissue regeneration. A. Quantification of mitotic numbers after disrupting ISC migration in intestines infected with *Ecc15* for 16hrs. B. Montage and quantification of ISC migratory behavior after modulating cell cycle progression. ISCs outlined in white, and montage tracks measured from the leading edge or, in cells that did not form protrusions, the most distal region of the cell cortex. Mock-treated and *Ecc15*-infected controls were taken from Figure 1B, as they served as genetic controls. C. Gut length decreases following 24hrs of *Ecc15* infection, but overcompensates after two days of recovery. Disrupting ISC migration impairs recovery. D. *P*.*e*. was fed to 10d females continuously. Disrupting ISC migration decreased survival rates. E. A model of non-canonical Wnt signaling-mediated regulation of ISC migration following injury. Mmp1 is activated in EE cells, which, in turn, facilitates activation of the non-canonical Wnt pathway, thereby promoting actin rearrangement. mean ± SEM (A) or mean ± SD (B,C); n≥16 flies (A), n≥5 flies (B), n≥8 flies (C), n≥44 flies (D); **P<0.01, ***P<0.001, based on one-way ANOVA with Tukey test (A,C), Student’s t-test (B), and log-rank test (D). Scale bar = 10µm. Timestamp indicated as hours:minutes. See also Supplemental Figure 7, Supplemental Video 24-25.

To further examine the relationship between cell migration and cell proliferation, we tested whether influencing cell cycle progression directly could also affect ISC migration. Indeed, overexpressing Wee1, a negative regulator of Cdk1^61^, impairs ISC migration after *Ecc15* infection (Fig. 6b and Supplemental Video 24). Conversely, accelerating G2 progression by depleting Wee1 was sufficient to induce ISC migration in undamaged tissue (Fig. 6b and Supplemental Video 25). This interdependency of cell cycle progression and cell migration in ISCs indicated that the induction of ISC migration is an integral part of the regenerative response of the intestinal epithelium. To further test this idea, we fed flies with *Ecc15* for 24 hours and then transferred them to normal food for 2 days to clear bacteria from the gut. The gut epithelium normally regenerates during this recovery phase, preserving gut size (Fig. 6c and Supplemental Fig. 7c). Depleting Dsh, Otk, or Klar did not affect gut size under homeostatic conditions (Supplemental Fig. 7d), but resulted in a reduced gut size after 2d recovery from *Ecc15* infection (Fig. 6c and Supplemental Fig. 7c). Impairing ISC migration still resulted in impaired regeneration after 1d recovery, but the effect was not as severe as gut size was still comparable between controls after *Ecc15* infection and 1d recovery (Supplemental Fig. 7d). We then investigated whether this impairment of regeneration also decreased survival rates after infection. Because *Ecc15* is not lethal to flies, we infected animals with *Pseudomonas Entomophila* (*P*.*e*.), an enteropathogen that induces tissue damage and causes death^62^, for 16 hours. Similar to *Ecc15, P*.*e*., induces migration (Supplemental Fig. 7e), and the depletion of Dsh, Otk, or Klar decreased survival rates after *P*.*e*. infection (Fig. 6d).

## Discussion

Our results uncover a mechanism by which EEs stimulate ISC migration towards an injured area in the intestinal epithelium (Fig. 6e). Injury induces Mmp expression in neighboring EEs, resulting in cleavage and release of the N-terminal domain of Otk. This, in turn, induces protrusion formation and migration in surrounding ISCs by activating Fz2/Dsh signaling, which promotes Rho/Rac-induced lamellipodia formation and Klar-mediated nuclear migration. We propose that the ensuing attraction of ISCs to the site of damage positions them to regenerate missing enterocytes *in situ*, and our data suggest that this is a critical component of a successful regenerative response in the intestinal epithelium. We further find that ISC migration is coupled to cell cycle progression in ISCs, indicating that the spatial reorganization of cells in the damaged epithelium may also be a prerequisite to the generation of new cells for epithelial replacement.

### Otk-dependent mechanisms regulate directed ISC migration after injury

Our study employs two experimental paradigms to probe ISC migration after epithelial injury: oral enteropathogen infection and localized damage using laser ablation. The epithelial injury induced by *Ecc15* infection induces a robust migratory response by ISCs, but lacks regional specificity due to the high titers of bacteria used in a laboratory setting. This is likely different in the wild, where lower titers of enteropathogens ingested with the food are likely to cause more localized damage. In our study we sought to mimic such localized damage using laser ablation, which induced migration of ISCs in close vicinity to the injury only, but allowed detailed characterization of directionality of protrusion formation and migration. A caveat of this approach is that the cultured and laser-injured intestines started to deteriorate after 2.5 hours of imaging, limiting the ability to track individual ISCs over time. To measure directionality of movement, we therefore had to rely on snapshots of the gut immediately after injury and 4.5 hours after ablation, limiting phototoxicity. These experiments revealed that ISCs extended protrusions towards the wound, and migrated towards and accumulated around the wound. However, they could not entirely rule out the possibility that the direction of ISC movement may be stochastic, and that ISCs pause or stop in the vicinity of the wound, resulting in the observed accumulation of ISCs around the wound.

Our data suggest that Wg secreted from EBs and Mmp-mediated release of N-terminal Otk from EEs are critical signals to initiate migration of ISCs after injury, and these secreted molecules may also serve as cues for chemotaxis. Metalloproteases play important roles during cell migration and chemotaxis^63,64^, and increased Mmp1 activity in EEs after injury may also facilitate the release of other signaling molecules. Consistent with a critical role for EE-derived signals, EEs have previously been reported to play important roles in regulating ISC proliferation^53^. Future studies are needed to identify possible additional chemoattractants, as well as to explore other downstream transduction pathways in migrating ISCs.

The exact molecular mechanism by which EE-derived N-terminal Otk induces ISC migration, however, remains unclear. Otk has been reported to be able to homodimerize^33^, and EE-derived Otk may interact with full-length Otk expressed in ISCs to stimulate non-canonical Wnt signaling. As proposed in other settings, Otk may function as co-receptor of Fz for Wnt ligands, stimulating non-canonical Wnt signaling and potentially inhibiting canonical Wnt signaling^27,33^. Importantly, our data suggest that activation of Dsh-mediated Wnt signaling is not required in EEs for the induction of ISC migration, and that the induction of Mmp and the cleavage of Otk are events occurring only in EEs, supporting a model in which EEs sense epithelial injury through an as yet to be identified mechanism, and then attract ISCs to the site of injury through secretion of N-terminal Otk.

In this model, EEs coordinate the directionality of ISC migration, while EBs support migration by serving as a major source of Wnt ligand. EB-derived Wg has been previously reported to be essential for ISC proliferation during regeneration^46^, and our data suggest that it is also required for effective ISC migration. It remains unclear, however, whether EBs serve as the only source of Wg for this process. Both Wnt2 and Wnt4 have been reported to interact with Otk *in vitro*^*27,33*^, and these other members of the Wnt family may also be secreted within the *Drosophila* intestine by other cell types to activate non-canonical Wnt signaling in ISCs.

### Interdependency of ISC migration and cell cycle progression

We find that ISC migration and proliferation are closely linked, and that modulating one of these processes affects the other. Because stem cells must divide at the right time and location to ensure proper regeneration, the coordination of migration and division may prevent ISCs from dividing prematurely before migrating closer to the wound. Migration serves to properly space cells within a tissue, and the increased ISC migration after induction of cell cycle progression may also ensure that newly formed cells are properly dispersed in the epithelium. The timing and regulation of this coordination, i.e. whether spatial cues trigger division when cells have reached their destination or whether the cytoskeletal rearrangements triggered during migration invariably induce cell cycle progression, will be fascinating to explore in future studies. Such coordination has been observed during development and in cancer cells through regulation of adhesion signaling by cell cycle-dependent kinases^65-67^. Exploring the mechanisms coordinating cell cycle and migration in adult stem cells further will deepen our understanding of the regenerative process.

### Adult stem cell migration in mammalian tissue

Our data reveal that stem cell migration plays a critical role in effective regeneration of the adult *Drosophila* intestine. It will be interesting to examine the extent of stem cell migration in adult mammalian tissue, and to determine whether the regulatory mechanisms observed in flies are conserved in these tissues. Migration of stem cells in adult mammalian tissue has largely been centered on hair follicle regeneration of the skin due to its accessibility. While canonical Wnt signaling is required for *de novo* hair follicle regeneration after wounding, the role of non-canonical Wnt signaling in the wound-induced migration of SCs remains to be characterized^1-3^,^4-6^. Differentiated epithelial cells within the mammalian intestine have recently been reported to actively migrate from the crypt to the top of the villus during steady-state tissue turnover^68^, but whether and how non-canonical Wnt signaling mediates cell migration in the mammalian intestine remains to be established. A role for Wnt5a (a ligand triggering non-canonical Wnt signaling) has been reported to promote homeostasis of the injured colonic epithelium, but it remains unclear whether this phenotype is associated with cell motility^69^. However, the organization of mammalian intestines into specialized regions, with ISCs sequestered at the base of crypts, make the tissue a poor analogue to study migration of stem cells. In contrast, other barrier tissues, such as the airway epithelium, share similar tissue organization and composition with the fly intestine. The basal stem cells in the mouse trachea, in particular, exhibit many conserved regulatory mechanisms with fly^70^. Exploring the conservation of role and regulation of stem cell migration in these mammalian tissues will improve our understanding of the regenerative response, and will likely lead to identification of novel targets and strategies for intervention for a wide range of human degenerative diseases.

## Supporting information

Supplemental Table 1 - List of Genotypes

Supplemental Video 1 Control Mock

Supplemental Video 2 Control Ecc15

Supplemental Video 3 EB Ecc15

Supplemental Video 4 EE Ecc15

Supplemental Video 5 Undamaged

Supplemental Video 6 Laser Ablation

Supplemental Video 7 LifeAct-GFP Mock

Supplemental Video 8 LifeAct-GFP Ecc15

Supplemental Video 9 LifeAct-GFP Ecc15 Cytochalasin B

Supplemental Video 10 Arp3 RNAi Ecc15

Supplemental Video 11 Klar RNAi Ecc15

Supplemental Video 12 Dsh RNAi Ecc15

Supplemental Video 13 Otk RNAi Ecc15

Supplemental Video 14 Dsh RNAi Ablation

Supplemental Video 15 Otk RNAi Ablation

Supplemental Video 16 LGK974 Ablation

Supplemental Video 17 Otk OE

Supplemental Video 18 EE>Otk OE

Supplemental Video 19 EE>Otk RNAi Ecc15

Supplemental Video 20 Ptk7

Supplemental Video 21 Otk RNAi Ptk7

Supplemental Video 22 EE>Timp OE Ecc15

Supplemental Video 23 TAPI-1 Ablation

Supplemental Video 24 Wee1 OE Ecc15

Supplemental Video 24 Wee1 RNAi

## Acknowledgement

We thank Drs. Jerome Korzelius, Lucy O’Brien, François Schweisguth and Andreas Wodarz for flies, Drs. Meredith Sagolla and Sarah Gierke for microscopy assistance, Dr. Scott Stawicki and Genentech, Inc. Protein Sciences for antibody and recombinant protein generation, and Xiaoyu Cai and Jesse Simons for technical assistance.

## Author Contributions

DJH, JY, and HJ conceived the project. DJH and HJ wrote the manuscript; JE helped established the laser ablation platform on the 2-photon microscopy system. DJH performed experiments and analyzed data. All authors read and approved the final manuscript.

## Methods

### Lead contact

Further information and requests for resources and reagents should be directed to and will be fulfilled by the Lead Contact, Heinrich Jasper. Unique biological materials generated in this study are readily available upon request, but will require a MTA.

### Experimental model and subject details

#### Drosophila stocks, culture, and husbandry

The following lines were obtained from the Bloomington *Drosophila* Stock Center: *UAS::nls-mCherry* (38425), *UAS::mCherry*^*RNAi*^ (35785), *UAS::GFP* (5431), *UAS::CD8-RFP* (27391), *UAS::dsh*^*RNAi*^ (31306), *UAS::eb1-GFP* (35512), *UAS::LifeAct-GFP* (35544), *how::gal4 (*1767), *UAS::klar*^*RNAi*^ (36721)^71^, *UAS::klar*^*RNAi*^ (28313), *UAS::otk*^*RNAi*^(25790), *UAS::otk2*^*RNAi*^ (57040), *UAS:: dsh*^*myc*^ (9453), *dsh*^*myc*^ (25385), *UAS::timp* (58708), *UAS::mmp1*^*RNAi*^ (31489)^72^, mmp1^*k04809*^*-lacZ* (12205), *UAS::wee1*^*RNAi*^ (55140)^73^, and *UAS::wee1-VFP* (65390). The following lines were obtained from the Vienna *Drosophila* Stock Center: *UAS::dia*^*RNAi*^ (v20518)^74^, *UAS::arp3*^*RNAi*^ (v108951)^75^, *UAS::rho1*^*RNAi*^ (v12734)^74^, *UAS::rac1*^*RNAi*^ (v49246)^76^, *UAS::dsh*^*RNAi*^ (v101525)^77^, *UAS::otk*^*RNAi*^ (v30833), *UAS::otk2*^*RNAi*^ (v106266), *UAS::fz*^*RNAi*^ (v43075)^78^, *UAS::fz2*^*RNAi*^ (v108988)^79^, *UAS::wg*^*RNAi*^ (v13352)^80^, *UAS::pan*^*RNAi*^ (v3014), and *UAS::arm*^*RNAi*^(v7767)^81^. The following lines were gifts: *esg::gal4, UAS::2xeYFP; Su(H)::Gal80, tub::Gal80*^*ts*82^, *mira-GFP* (Dr. François Schweisguth)^54^, *UAS::otk-GFP* (Dr. Andreas Wodarz)^33^, *tub::Gal80*^*ts*^; *pros::Gal4* (Dr. Jerome Korzelius)^83^, *mex1-gal4* (Dr. Lucy O’Brien)^15^, *esg::FlipOut*^60^, and *Su(H)GBE::Gal4*^84^. The genotype of flies used in each figure is detailed in Supplemental Table 1. All RNAi lines except for Dsh (31306), Klar (28313), Otk, and Otk2 were previously validated, as referenced, by either qPCR or immunofluorescence. Two different RNAi lines were used for Dsh, Otk, Otk2, and Klar to minimize off-target effects. Lines 25790, 28313, and 57040 were only used for Supplemental Fig. 4b. Line 31306 was used for Fig. 3c, Supplemental Fig. 4b, and Supplemental 5d.

Flies were raised at 18°C to prevent RNAi or protein expression during development and 65% humidity, on a 12-hour light/dark cycle. To induce genetic expression, flies were shifted to 29°C for three days prior to bacterial infection or mock-treatment (for a total of ∼four days induction). Otherwise, flies were shifted from 18°C to 29°C for four days. Standard fly food was prepared with the following recipe: 1 L distilled water, 7.3 ml EtOH, 13 g agar, 22g molasses, 65g malt extract, 18g brewer’s yeast, 80g corn flour, 10g soy flour, 6.2ml propionic acid, and 2g methyl-p-benzoate. Flies were manipulated by CO_2_ anesthesia. Only female flies were used in this study because females exhibit higher intestinal turnover rates than males. All flies were dissected at 10d except for gut size experiments, which were dissected at 11d.

### Method details

#### Drosophila Ecc15 and P.e. infection

For *Ecc15* infection, *Ecc15* was cultured in 15ml of LB medium for 16 hours at 30°. The culture was pelleted, resuspended in 400µL of 5% sucrose in water, and fed to two-hour starved flies on Whatman paper for 16 hours. Intestines were subsequently dissected. For gut size experiments, flies were fed with *Ecc15* for 24 hours before transferred to standard fly food for up to 48 hours. For *P*.*e*. infection, *P*.*e*. was cultured in 20ml of LB medium for 16 hours at 30°. The culture was pelleted, resuspended in 400µL of 5% sucrose in water, and fed to two-hour starved flies for either 16 hours (for live imaging experiments) or the duration of the experiment (for survival experiments). For survival experiments, 40-60 10d flies were housed together and continuously fed with *P*.*e*, with 300µl of 5% sucrose in water added daily. Dead flies were counted daily. For mock controls, flies were starved for two hours and fed 5% sucrose on a vial containing Whatman filter paper for 16 hours. Survival experiments were repeated three times.

#### Live imaging of *Drosophila* intestines

Intestines from adult female flies were dissected in culture media containing 2mM CaCl_2_, 5mM KCl, 5mM HEPES, 8.2mM MgCl_2_, 108mM NaCl, 4mM NaHCO_3_, NaH_2_PO_4_, 10mM sucrose, 5mM trehalose, and 2% fetal bovine serum (Adult Hemolymph-like Saline, AHLS). Intestines were transferred to a 35mm glass bottom dish (MatTek, P35G-1.5-14-C), embedded in 3% low melting agarose (in AHLS), and submerged in AHLS. The posterior midgut was imaged at intervals of 10 min for up to three hours on a Yokogawa CSU-W1/Zeiss 3i Marianas spinning disk confocal microscopy system with a 40x PlanFleur objective or a Andor CSU-X1/Nikon Ti-E spinning disk confocal microscopy system with a 40x PlanFleur objective, or, for laser ablation experiments, on a Bruker Ultima In Vivo Multiphoton/Newport MaiTai DeepSee Spectra Physics 2-photon microscopy system with a 20x Olympus 1.0 water immersion objective.

#### Laser ablation of *Drosophila* intestines

Ablated intestines and corresponding undamaged controls were imaged with a 960nm wavelength, 2.8µs dwell time, and ∼25mW power, using a Bruker Ultima In Vivo Multiphoton/Newport MaiTai DeepSee Spectra Physics 2-photon microscopy system with a 20x Olympus 1.0 water immersion objective. A wound of a ∼30µm diameter was created in the posterior midgut with a 880nm wavelength, 20µs dwell time, ∼278mW power, and 60x zoom on the same system. Ablated guts were live imaged on the 2-photon system at intervals of 10 min for ∼2.5hrs. For fixed analysis, ablated guts were cultured in AHLS for either 1hr or 4.5hrs prior to fixation.

#### Treatment of *Drosophila* intestine with small molecule inhibitors and recombinant protein

Intestines from adult female flies were dissected in AHLS (see Live Imaging of *Drosophila* intestines) and treated with small molecule inhibitors or recombinant protein in AHLS 30 minutes prior to imaging and throughout the live imaging process. Mock-treated intestines were incubated in 0.1% DMSO in AHLS 30 minutes prior to imaging and throughout the live imaging process.

Small molecule inhibitors and concentrations used in this study: Blebbistatin (20µM, Sigma Aldrich, B0560), Cytochalasin B (10µM, Sigma Aldrich, C6762), LGK974 (1µM, Millipore Sigma Chemicals, 531091), and TAPI-1 (20µM, EMD Millipore, 59053). Recombinant protein and concentrations used in this study: C-terminal Ptk7 (1µg/ml, see Recombinant C-terminal Ptk7 generation).

#### Immunostaining of *Drosophila* intestine

Intestines from adult female flies were dissected in PBS (Phosphate-buffered saline, pH 7.4), and fixed for 25 minutes at room temperature in media containing 100mM glutamic acid, 25mM KCl, 20mM MgSO_4_, 4mM sodium phosphate, 1mM MgCl_2_, and 4% formaldehyde. Intestines were washed with PBS and stained in PBS, 0.3% Triton X-100 supplemented with 5% donkey serum. Intestines were incubated in primary antibody overnight at 4°C antibody, washed in PBS +0.01% Tween-20, and incubated in secondary antibodies and Hoechst stain (1:1000) for 2 hours at room temperature.

Antibodies used in this study: Mouse monoclonal against Prospero (1:300, DHSB, MR1A, RRID:AB_528440), Myc (1:300, Abcam, ab32, RRID:AB_303599), and Mmp1 (1:10, DHSB, 5H7B11, RRID: AB_579779; 3A6B4, RRID:AB_579780; 3B8D12, RRID: AB_579781); rabbit polyclonal against ß-galactosidase (1:300, MP Biomedicals, 085597-CF), phospho-Histone H3 (1:1000, EMD Millipore, 06-570, RRID:AB_310177), and Otk (1:50, see Otk antibody generation); and chicken polyclonal against GFP (1:300, Abcam, ab13970, RRID:AB_300798). Secondary antibodies were cyanine dyes from Jackson ImmunoResearch Laboratories (1:300). Anti-GFP antibodies were used to amplify the signal in cells overexpressing GFP-tagged Otk. For Mmp1 staining, all three Mmp1 antibodies were used in conjunction at 1:10 in a 1:1:1 mixture as previously reported^85^.

#### Otk antibody and recombinant Ptk7 generation

To generate polyclonal Otk antibodies, rabbits were immunized against immunogens consisting of amino acids 23-581 for recognition of the extracellular domain or 632-1033 for recognition of the intracellular domain (Genentech, Inc. Protein Sciences). Sera were extracted and purified using Protein A, resulting in IgG polysera. Reactivity was confirmed by immunofluorescence. To generate recombinant murine extracellular Ptk7, Met1-Glu683 was cloned into a modified pRK vector containing a CMV promoter and C-terminal Flag tag (Genentech, Inc. Protein Sciences). Protein was expressed in HEK293 cells using standard transfection protocols and supernatant was harvested after seven days. Clarified supernatant was passed over Flag-resin equilibrated with buffer A (25 mM Tris pH 7.5, 150 mM NaCl, 5 mM EDTA), washed with buffer B (25 mM Tris pH 7.5, 150 mM NaCl, 5 mM EDTA, 0.2% Triton X-114, 0.2% Triton X100), eluted in buffer C (50 mM sodium citrate, 150 mM NaCl, pH 3.0), and neutralized by addition of 1 M Tris pH 8.0. Protein was concentrated and passed over a Superdex 200 column in phosphate buffered saline, and the peak fraction was collected. SDS-PAGE and mass spectrometry (ISD MALDI) were used to confirm high purify and identity of the protein (starting at residue Ala23).

#### Esg::FlipOut induction

Flies were raised at 18°C and shifted to 29°C for three days starting at 6d. Flies were then fed *Ecc15* in 5% sucrose or 5% sucrose alone at 9d and maintained at 29°C for 24hrs before dissected. The number of cells per clone was quantified.

#### Microscopy and image analysis

Images of fixed tissue were taken either with a Yokogawa CSU-W1/Zeiss 3i Marianas spinning disk confocal microscopy system or a Andor CSU-X1/Nikon Ti-E spinning disk confocal microscopy system. Intestines were imaged using a 40x PlanFleur objective or, for gut size measurements, a 10x PlanFleur objective. Images were analyzed and processed using ImageJ (NIH, Bethesda, MD) and Adobe Photoshop. Figures were composed in Adobe Illustrator. Except for quantification of PH3+ cells and gut length, which was performed in the entire gut, only the R4 region of the posterior midgut was analyzed for consistency.

### Quantification and statistical analysis

All quantifications were performed manually using ImageJ software (NIH, Bethesda, MD). Except for quantification of PH3+ cells and gut length, which was performed in the entire gut, only the R4 region of the posterior midgut was analyzed for consistency.

#### Quantification of gut size and immunostaining

Intestines were dissected from females at 10d of age, except for gut length analyses, which were at 11d. Mitotic numbers were counted from the entire gut as determined by phospho-histone H3 positivity. Gut length was measured from the beginning of the beginning of the anterior midgut to the end of the posterior midgut. The presence of cortical Dsh was identified by localization of Dsh around the periphery of the ISC, as determined by cytoplasmic GFP expression. Cell-type specific expression of Otk and Mmp1 was determined by ISC- and EE-specific expression of GFP, or prospero immunostaining for EEs. Dsh and Mmp1 expression in ablated intestines was only quantified within a 100µm radius of the ablation site. For measurement of N-terminal Otk and Mmp1 intensity, fluorescence levels were set to the same baseline across all images. Relative N-terminal Otk levels were measured by normalizing the corrected total cell fluorescence (CTCF) of N-terminal Otk in the cell outline with the CTCF of C-terminally GFP-tagged Otk. Relative Mmp1 levels were measured by normalizing mean fluorescence of Mmp1 in the cell outline with the mean fluorescence of the same area in adjacent regions.

#### Analysis of intestinal stem cell migration

Only the R4 region of the posterior midgut, chosen randomly, was imaged. Quantification of migratory behavior consisted of assaying for protrusion formation and cell migration, and were only performed on guts with minimal sample movement as determined by auto fluorescence of the tissue. ISCs were scored as exhibiting protrusion formation if a protrusion was extended at any point during a 2.5hr live imaging duration. ISCs were scored as exhibiting cell migration if the cell body translocated at least 3µm during a 2.5hr live imaging duration, using the tissue background as a reference point, if necessary. Tracks were measured every ten minutes using the Manual Tracking plugin (Fabrice Cordeli, Curie Institute, Paris, France) in ImageJ (NIH, Bethesda, MD), tracing either the edge of the protrusion or the center of the cell body. The starting point of each ISC was designated as the center of the Cartesian plane (0,0), and the x,y coordinate of the ISC at each time point was normalized to its starting point. For ISCs that did not exhibit protrusion formation, the most distal region of the cell cortex was tracked instead. For laser ablation experiments, ISC migratory behavior was assayed within 75µms from the site of injury.

#### Analysis of migration directionality

To assay ISC accumulation around the periphery of the wound in fixed, ablated intestines (Supplemental 2c), the ISC numbers within a 40µm radius of the ablation site were counted and compared with the ISC numbers within the same area of the contralateral side of the intestine. ISCs within the ablation site area on the contralateral side were not counted. For undamaged controls, ISCs were counted on contralateral sides of the intestine in a 200µm × 200µm area. To quantify the angle of ISC protrusions in relation to the wound of an ablated gut (Supplemental Fig 2a), the angle was measured between the vector of the shortest distance from the center of the cell body to the periphery of the wound and the vector from the center of the cell body to the center of the leading edge. To plot ISC position in ablated tissue (Supplemental 2b), the periphery of the wound closest to a given ISC was designated as the center of the Cartesian plane (0,0). The starting position of the ISC was the x,y coordinate of the cell cortex closest to the wound immediately after ablation, and normalized to the x,y coordinate of the wound periphery closest to the ISC. The ending position of the ISC was the x,y coordinate of the cell cortex closest to the wound 4.5hrs post ablation, and normalized to the x,y coordinate of the wound periphery closest to the ISC. For example, if the x,y coordinate of an ISC was measured at 108, 148 and the wound was measured at 133, 148, the wound would be set at 0,0 and the normalized x,y coordinate of the ISC would be -25, 0. In control, unablated tissue, a single arbitrary point in the middle of the gut was designated as the center of the Cartesian plane (0,0). ISC position was determined by the x,y coordinate of the cell cortex closest to the designated point, and normalized to the x,y coordinate of the designated point. Change in displacement was calculated from the difference in distance between the starting ISC position and the Cartesian origin, and the ending ISC position and the Cartesian origin.

#### Statistical analysis

Each sample, ‘n’, is from at least two independent experiments, and is defined in the Figure Legends depending on experiment. Additional statistical details for each experiment are also noted in the Figure Legends. Statistical analyses were performed with Prism (GraphPad Software, La Jolla, CA, USA). A Student’s t-test was used to determine statistical significance between two independent groups. A one-way ANOVA with Tukey test was used to determine statistical significance with multiple comparisons between three or more independent groups. A chi-square test was used to test for statistical significance when comparing data according to a set hypothesis (percentage of protrusions formed towards versus away from wound). A log-rank test was used to test for statistical significance in *P*.*e*. survival assays. Significance was accepted at the level of p < 0.05. No statistical methods were used to predetermine sample sizes, but our sample sizes are similar to those generally employed in the field. The specific region imaged was chosen at random within the R4 of the posterior midgut. All experiments (from both live and fixed samples) were quantified blindly.

## Data and code availability

This study did not generate any data sets or code.

## References

1 Dekoninck, S. & Blanpain, C. Stem cell dynamics, migration and plasticity during wound healing. Nature cell biology 21, 18–24, doi:10.1038/s41556-018-0237-6 (2019).

2 de Lucas, B., Pérez, L. M. & Gálvez, B. G. Importance and regulation of adult stem cell migration. 22, 746–754, doi:10.1111/jcmm.13422 (2018).

3 Fu, X. et al. Mesenchymal Stem Cell Migration and Tissue Repair. Cells 8, 784, doi:10.3390/cells8080784 (2019).

4 Wu, X. et al. Skin stem cells orchestrate directional migration by regulating microtubule-ACF7 connections through GSK3β. Cell 144, 341–352, doi:10.1016/j.cell.2010.12.033 (2011).

5 Park, S. et al. Tissue-scale coordination of cellular behaviour promotes epidermal wound repair in live mice. Nature cell biology 19, 155–163, doi:10.1038/ncb3472 (2017).

6 Aragona, M. et al. Defining stem cell dynamics and migration during wound healing in mouse skin epidermis. Nature communications 8, 14684, doi:10.1038/ncomms14684 (2017).

7 Wright, D. E., Wagers, A. J., Gulati, A. P., Johnson, F. L. & Weissman, I. L. Physiological migration of hematopoietic stem and progenitor cells. Science 294, 1933–1936, doi:10.1126/science.1064081 (2001).

8 Tang, Y. et al. TGF-β1–induced migration of bone mesenchymal stem cells couples bone resorption with formation. Nature Medicine 15, 757–765, doi:10.1038/nm.1979 (2009).

9 Lin, W. et al. Mesenchymal stem cells homing to improve bone healing. Journal of orthopaedic translation 9, 19–27, doi:10.1016/j.jot.2017.03.002 (2017).

10 Sahin, A. O. & Buitenhuis, M. Molecular mechanisms underlying adhesion and migration of hematopoietic stem cells. Cell Adh Migr 6, 39–48, doi:10.4161/cam.18975 (2012).

11 Hu, D. J. & Jasper, H. Control of Intestinal Cell Fate by Dynamic Mitotic Spindle Repositioning Influences Epithelial Homeostasis and Longevity. Cell reports 28, 2807–2823.e2805, doi:10.1016/j.celrep.2019.08.014 (2019).

12 Morris, O., Deng, H., Tam, C. & Jasper, H. Warburg-like Metabolic Reprogramming in Aging Intestinal Stem Cells Contributes to Tissue Hyperplasia. Cell reports 33, 108423, doi:10.1016/j.celrep.2020.108423 (2020).

13 Ohlstein, B. & Spradling, A. The adult Drosophila posterior midgut is maintained by pluripotent stem cells. Nature 439, 470–474 (2006).

14 Martin, J. L. et al. Long-term live imaging of the Drosophila adult midgut reveals real-time dynamics of division, differentiation and loss. eLife 7, e36248, doi:10.7554/eLife.36248 (2018).

15 Koyama, L. A. J. et al. Bellymount enables longitudinal, intravital imaging of abdominal organs and the gut microbiota in adult Drosophila. PLOS Biology 18, e3000567, doi:10.1371/journal.pbio.3000567 (2020).

16 Lee, J., Cabrera, A. J. H., Nguyen, C. M. T. & Kwon, Y. V. Dissemination of Ras(V12)-transformed cells requires the mechanosensitive channel Piezo. Nature communications 11, 3568–3568, doi:10.1038/s41467-020-17341-y (2020).

17 Micchelli, C. A. & Perrimon, N. Evidence that stem cells reside in the adult Drosophila midgut epithelium. Nature 439, 475–479, doi:http://www.nature.com/nature/journal/v439/n7075/suppinfo/nature04371_S1.html (2006).

18 Buchon, N., Broderick, N. A., Chakrabarti, S. & Lemaitre, B. Invasive and indigenous microbiota impact intestinal stem cell activity through multiple pathways in Drosophila. Genes & development 23, 2333–2344, doi:10.1101/gad.1827009 (2009).

19 O’Brien, L. E., Soliman, S. S., Li, X. & Bilder, D. Altered modes of stem cell division drive adaptive intestinal growth. Cell 147, 603–614, doi:10.1016/j.cell.2011.08.048 (2011).

20 Mayor, R. & Etienne-Manneville, S. The front and rear of collective cell migration. Nat Rev Mol Cell Biol 17, 97–109, doi:10.1038/nrm.2015.14 (2016).

21 Pollard, T. D. & Cooper, J. A. Actin, a central player in cell shape and movement. Science 326, 1208–1212, doi:10.1126/science.1175862 (2009).

22 Watanabe, N., Kato, T., Fujita, A., Ishizaki, T. & Narumiya, S. Cooperation between mDia1 and ROCK in Rho-induced actin reorganization. Nature cell biology 1, 136–143, doi:10.1038/11056 (1999).

23 Nobes, C. D. & Hall, A. Rho GTPases control polarity, protrusion, and adhesion during cell movement. J Cell Biol 144, 1235–1244, doi:10.1083/jcb.144.6.1235 (1999).

24 Michaelis, U. R. Mechanisms of endothelial cell migration. Cellular and molecular life sciences : CMLS 71, 4131–4148, doi:10.1007/s00018-014-1678-0 (2014).

25 Ridley, A. J. et al. Cell Migration: Integrating Signals from Front to Back. Science 302, 1704, doi:10.1126/science.1092053 (2003).

26 Vicente-Manzanares, M., Ma, X., Adelstein, R. S. & Horwitz, A. R. Non-muscle myosin II takes centre stage in cell adhesion and migration. Nat Rev Mol Cell Biol 10, 778–790, doi:10.1038/nrm2786 (2009).

27 Peradziryi, H. et al. PTK7/Otk interacts with Wnts and inhibits canonical Wnt signalling. The EMBO journal 30, 3729–3740, doi:https://doi.org/10.1038/emboj.2011.236 (2011).

28 Davey, C. F. & Moens, C. B. Planar cell polarity in moving cells: think globally, act locally. Development 144, 187, doi:10.1242/dev.122804 (2017).

29 Lu, X. et al. PTK7/CCK-4 is a novel regulator of planar cell polarity in vertebrates. Nature 430, 93–98, doi:10.1038/nature02677 (2004).

30 Martinez, S. et al. The PTK7 and ROR2 Protein Receptors Interact in the Vertebrate WNT/Planar Cell Polarity (PCP) Pathway. The Journal of biological chemistry 290, 30562–30572, doi:10.1074/jbc.M115.697615 (2015).

31 Yen, W. W. et al. PTK7 is essential for polarized cell motility and convergent extension during mouse gastrulation. Development (Cambridge, England) 136, 2039–2048, doi:10.1242/dev.030601 (2009).

32 De Calisto, J., Araya, C., Marchant, L., Riaz, C. F. & Mayor, R. Essential role of non-canonical Wnt signalling in neural crest migration. Development 132, 2587, doi:10.1242/dev.01857 (2005).

33 Linnemannstöns, K. et al. The PTK7-related transmembrane proteins off-track and off-track 2 are co-receptors for Drosophila Wnt2 required for male fertility. PLoS Genet 10, e1004443–e1004443, doi:10.1371/journal.pgen.1004443 (2014).

34 Gao, C. & Chen, Y. G. Dishevelled: The hub of Wnt signaling. Cellular signalling 22, 717–727, doi:10.1016/j.cellsig.2009.11.021 (2010).

35 Shnitsar, I. & Borchers, A. PTK7 recruits dsh to regulate neural crest migration. Development 135, 4015–4024, doi:10.1242/dev.023556 (2008).

36 Gómez-Orte, E., Sáenz-Narciso, B., Moreno, S. & Cabello, J. Multiple functions of the noncanonical Wnt pathway. Trends in genetics : TIG 29, 545–553, doi:10.1016/j.tig.2013.06.003 (2013).

37 McGuire, S. E., Mao, Z. & Davis, R. L. Spatiotemporal gene expression targeting with the TARGET and gene-switch systems in Drosophila. Science’s STKE : signal transduction knowledge environment 2004, pl6, doi:10.1126/stke.2202004pl6 (2004).

38 Wang, L., Karpac, J. & Jasper, H. Promoting longevity by maintaining metabolic and proliferative homeostasis. J Exp Biol 217, 109–118, doi:10.1242/jeb.089920 (2014).

39 Buchon, N., Broderick, N. A., Poidevin, M., Pradervand, S. & Lemaitre, B. Drosophila Intestinal Response to Bacterial Infection: Activation of Host Defense and Stem Cell Proliferation. Cell Host & Microbe 5, 200–211, doi:https://doi.org/10.1016/j.chom.2009.01.003 (2009).

40 Ayyaz, A. & Jasper, H. Intestinal inflammation and stem cell homeostasis in aging Drosophila melanogaster. Frontiers in cellular and infection microbiology 3, 98, doi:10.3389/fcimb.2013.00098 (2013).

41 MacLean-Fletcher, S. & Pollard, T. D. Mechanism of action of cytochalasin B on actin. Cell 20, 329–341, doi:https://doi.org/10.1016/0092-8674(80)90619-4 (1980).

42 Straight, A. F. et al. Dissecting Temporal and Spatial Control of Cytokinesis with a Myosin II Inhibitor. Science 299, 1743, doi:10.1126/science.1081412 (2003).

43 Krause, M. & Gautreau, A. Steering cell migration: lamellipodium dynamics and the regulation of directional persistence. Nat Rev Mol Cell Biol 15, 577–590, doi:10.1038/nrm3861 (2014).

44 Starr, D. A. & Fridolfsson, H. N. in Annual Review of Cell and Developmental Biology Vol. 26 421–444 (2010).

45 Liu, J. et al. Targeting Wnt-driven cancer through the inhibition of Porcupine by LGK974. Proceedings of the National Academy of Sciences 110, 20224, doi:10.1073/pnas.1314239110 (2013).

46 Cordero, J. B., Stefanatos, R. K., Scopelliti, A., Vidal, M. & Sansom, O. J. Inducible progenitor-derived Wingless regulates adult midgut regeneration in Drosophila. The EMBO journal 31, 3901–3917, doi:10.1038/emboj.2012.248 (2012).

47 Brunner, E., Peter, O., Schweizer, L. & Basler, K. pangolinencodes a Lef-1 homologue that acts downstream of Armadillo to transduce the Wingless signal in Drosophila. Nature 385, 829–833, doi:10.1038/385829a0 (1997).

48 Clevers, H. & Nusse, R. Wnt/β-catenin signaling and disease. Cell 149, 1192–1205, doi:10.1016/j.cell.2012.05.012 (2012).

49 Axelrod, J. D., Miller, J. R., Shulman, J. M., Moon, R. T. & Perrimon, N. Differential recruitment of Dishevelled provides signaling specificity in the planar cell polarity and Wingless signaling pathways. Genes & development 12, 2610–2622, doi:10.1101/gad.12.16.2610 (1998).

50 Axelrod, J. D. Unipolar membrane association of Dishevelled mediates Frizzled planar cell polarity signaling. Genes & development 15, 1182–1187, doi:10.1101/gad.890501 (2001).

51 Park, T. J., Gray, R. S., Sato, A., Habas, R. & Wallingford, J. B. Subcellular localization and signaling properties of dishevelled in developing vertebrate embryos. Current biology : CB 15, 1039–1044, doi:10.1016/j.cub.2005.04.062 (2005).

52 Wang, Y. & Naturale, V. F. Planar Cell Polarity Effector Fritz Interacts with Dishevelled and Has Multiple Functions in Regulating PCP. 7, 1323–1337, doi:10.1534/g3.116.038695 (2017).

53 Amcheslavsky, A. et al. Enteroendocrine Cells Support Intestinal Stem-Cell-Mediated Homeostasis in Drosophila. Cell reports 9, 32–39, doi:https://doi.org/10.1016/j.celrep.2014.08.052 (2014).

54 Bardin, A. J., Perdigoto, C. N., Southall, T. D., Brand, A. H. & Schweisguth, F. Transcriptional control of stem cell maintenance in the Drosophila intestine. Development 137, 705–714, doi:10.1242/dev.039404 (2010).

55 Na, H.-W., Shin, W.-S., Ludwig, A. & Lee, S.-T. The cytosolic domain of protein-tyrosine kinase 7 (PTK7), generated from sequential cleavage by a disintegrin and metalloprotease 17 (ADAM17) and γ-secretase, enhances cell proliferation and migration in colon cancer cells. The Journal of biological chemistry 287, 25001–25009, doi:10.1074/jbc.M112.348904 (2012).

56 Golubkov, V. S. et al. The Wnt/planar cell polarity protein-tyrosine kinase-7 (PTK7) is a highly efficient proteolytic target of membrane type-1 matrix metalloproteinase: implications in cancer and embryogenesis. The Journal of biological chemistry 285, 35740–35749, doi:10.1074/jbc.M110.165159 (2010).

57 Golubkov, V. S. et al. Protein-tyrosine pseudokinase 7 (PTK7) directs cancer cell motility and metastasis. The Journal of biological chemistry 289, 24238–24249, doi:10.1074/jbc.M114.574459 (2014).

58 Azpurua, J., Mahoney, R. E. & Eaton, B. A. Transcriptomics of aged Drosophila motor neurons reveals a matrix metalloproteinase that impairs motor function. Aging cell 17, doi:10.1111/acel.12729 (2018).

59 Muliyil, S. et al. ADAM17-triggered TNF signalling protects the ageing Drosophila retina from lipid droplet-mediated degeneration. The EMBO journal 39, e104415–e104415, doi:10.15252/embj.2020104415 (2020).

60 Jiang, H. et al. Cytokine/Jak/Stat signaling mediates regeneration and homeostasis in the Drosophila midgut. Cell 137, 1343–1355, doi:10.1016/j.cell.2009.05.014 (2009).

61 Stumpff, J., Duncan, T., Homola, E., Campbell, S. D. & Su, T. T. Drosophila Wee1 kinase regulates Cdk1 and mitotic entry during embryogenesis. Current biology : CB 14, 2143–2148, doi:10.1016/j.cub.2004.11.050 (2004).

62 Vodovar, N. et al. *Drosophila* host defense after oral infection by an entomopathogenic *Pseudomonas* species. Proceedings of the National Academy of Sciences of the United States of America 102, 11414, doi:10.1073/pnas.0502240102 (2005).

63 Page-McCaw, A., Ewald, A. J. & Werb, Z. Matrix metalloproteinases and the regulation of tissue remodelling. Nature Reviews Molecular Cell Biology 8, 221–233, doi:10.1038/nrm2125 (2007).

64 Chen, P. & Parks, W. C. Role of matrix metalloproteinases in epithelial migration. J Cell Biochem 108, 1233–1243, doi:10.1002/jcb.22363 (2009).

65 Ridenour, D. A. et al. The neural crest cell cycle is related to phases of migration in the head. Development 141, 1095–1103, doi:10.1242/dev.098855 (2014).

66 Chu, T. L. H. et al. Cell Cycle-Dependent Tumor Engraftment and Migration Are Enabled by Aurora-A. Molecular cancer research : MCR 16, 16–31, doi:10.1158/1541-7786.mcr-17-0417 (2018).

67 Jones, M. C., Zha, J. & Humphries, M. J. Connections between the cell cycle, cell adhesion and the cytoskeleton. Philosophical transactions of the Royal Society of London. Series B, Biological sciences 374, 20180227, doi:10.1098/rstb.2018.0227 (2019).

68 Krndija, D. et al. Active cell migration is critical for steady-state epithelial turnover in the gut. Science 365, 705, doi:10.1126/science.aau3429 (2019).

69 Miyoshi, H., Ajima, R., Luo, C. T., Yamaguchi, T. P. & Stappenbeck, T. S. Wnt5a potentiates TGF-β signaling to promote colonic crypt regeneration after tissue injury. Science 338, 108–113, doi:10.1126/science.1223821 (2012).

70 Haller, S. et al. mTORC1 Activation during Repeated Regeneration Impairs Somatic Stem Cell Maintenance. Cell stem cell 21, 806–818.e805, doi:10.1016/j.stem.2017.11.008 (2017).

71 Collins, M. A. et al. Emery-Dreifuss muscular dystrophy-linked genes and centronuclear myopathy-linked genes regulate myonuclear movement by distinct mechanisms. Molecular biology of the cell 28, 2303–2317, doi:10.1091/mbc.E16-10-0721 (2017).

72 Sauerwald, J., Backer, W., Matzat, T., Schnorrer, F. & Luschnig, S. Matrix metalloproteinase 1 modulates invasive behavior of tracheal branches during entry into Drosophila flight muscles. eLife 8, e48857, doi:10.7554/eLife.48857 (2019).

73 Sopko, R. et al. Combining genetic perturbations and proteomics to examine kinase-phosphatase networks in Drosophila embryos. Dev Cell 31, 114–127, doi:10.1016/j.devcel.2014.07.027 (2014).

74 Widmann, T. J. & Dahmann, C. Dpp signaling promotes the cuboidal-to-columnar shape transition of Drosophila wing disc epithelia by regulating Rho1. Journal of cell science 122, 1362–1373, doi:10.1242/jcs.044271 (2009).

75 Verboon, J. M., Rahe, T. K., Rodriguez-Mesa, E. & Parkhurst, S. M. Wash functions downstream of Rho1 GTPase in a subset of Drosophila immune cell developmental migrations. Molecular biology of the cell 26, 1665–1674, doi:10.1091/mbc.E14-08-1266 (2015).

76 Baek, S. H., Kwon, Y. C., Lee, H. & Choe, K. M. Rho-family small GTPases are required for cell polarization and directional sensing in Drosophila wound healing. Biochemical and biophysical research communications 394, 488–492, doi:10.1016/j.bbrc.2010.02.124 (2010).

77 Agrawal, T. & Hasan, G. Maturation of a central brain flight circuit in Drosophila requires Fz2/Ca2+ signaling. Elife 4, doi:10.7554/eLife.07046 (2015).

78 Trujillo, G. V. et al. The canonical Wingless signaling pathway is required but not sufficient for inflow tract formation in the Drosophila melanogaster heart. Developmental biology 413, 16–25, doi:10.1016/j.ydbio.2016.03.013 (2016).

79 Chaudhary, V. et al. Robust Wnt signaling is maintained by a Wg protein gradient and Fz2 receptor activity in the developing Drosophila wing. 146, doi:10.1242/dev.174789 (2019).

80 Tan, Y., Yu, D., Busto, G. U., Wilson, C. & Davis, R. L. Wnt signaling is required for long-term memory formation. Cell reports 4, 1082–1089, doi:10.1016/j.celrep.2013.08.007 (2013).

81 Sarpal, R. et al. Mutational analysis supports a core role for Drosophila α-catenin in adherens junction function. Journal of cell science 125, 233–245, doi:10.1242/jcs.096644 (2012).

82 Wang, L., Zeng, X., Ryoo, H. D. & Jasper, H. Integration of UPRER and oxidative stress signaling in the control of intestinal stem cell proliferation. PLoS Genet 10, e1004568, doi:10.1371/journal.pgen.1004568 (2014).

83 Grosjean, Y., Balakireva, M., Dartevelle, L. & Ferveur, J. F. PGal4 excision reveals the pleiotropic effects of Voila, a Drosophila locus that affects development and courtship behaviour. Genetical research 77, 239–250, doi:10.1017/s0016672301005006 (2001).

84 Zeng, X., Chauhan, C. & Hou, S. X. Characterization of midgut stem cell-and enteroblast-specific Gal4 lines in drosophila. Genesis (New York, N.Y. : 2000) 48, 607–611, doi:10.1002/dvg.20661 (2010).

85 Lee, S. H., Park, J. S., Kim, Y. S., Chung, H. Y. & Yoo, M. A. Requirement of matrix metalloproteinase-1 for intestinal homeostasis in the adult Drosophila midgut. Exp Cell Res 318, 670–681, doi:10.1016/j.yexcr.2012.01.004 (2012).

