## Supplemental Figures, Supplemental Figure Legends, and Supplemental Video Legends for "Non-canonical Wnt signaling promotes directed migration of intestinal stem cells to sites of injury"

### Supplemental Figure 1

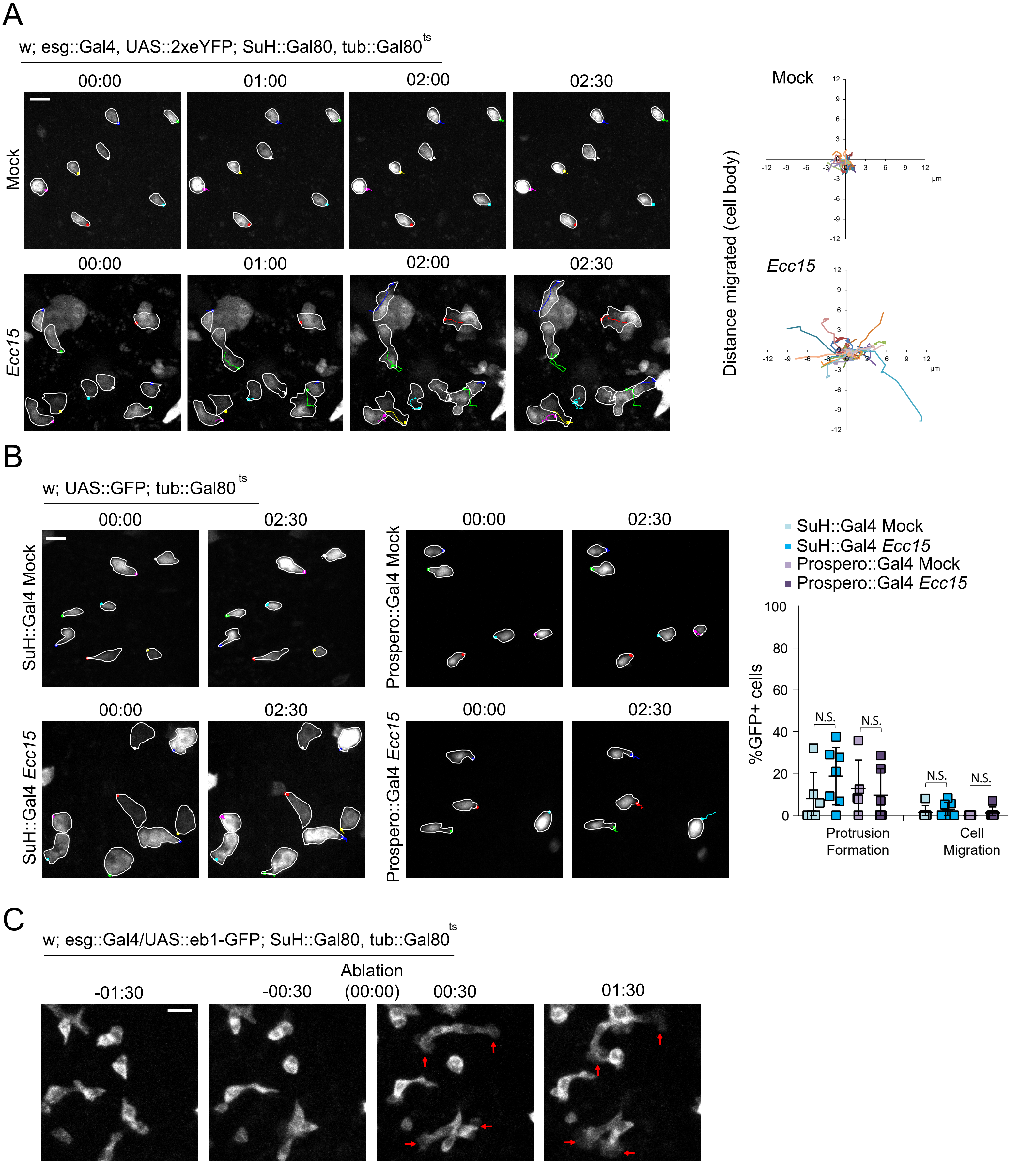

**Supplemental Figure 1: Additional characterization of cell migration in *Drosophila* intestine, Related to Figure 1.**

A. Montage of ISC migration and tracings of the displacement of the ISC body in mock-treated and *Ecc15*-infected intestines. ISCs outlined in white, and montage tracks measured from the leading edge or, in cells that did not form protrusions, the most distal region of the cell cortex. Starting x,y coordinate of ISCs normalized to 0,0. B. Montage and quantification of EB and EE migratory behavior in mock-treated and *Ecc15*-infected intestines. GFP+ cells outlined in white, and montage tracks measured from the most distal region of the cell cortex. C. Montage of cytoskeleton dynamics pre- and post-ablation. Ablation occurred at time point 00:00. Red arrows indicate protrusions. mean ± SD; n≥5 flies (B); N.S. = not significant, based on Student's t-test. Scale bar = 10µm. Timestamp indicated as hours:minutes.

### Supplemental Figure 2

A

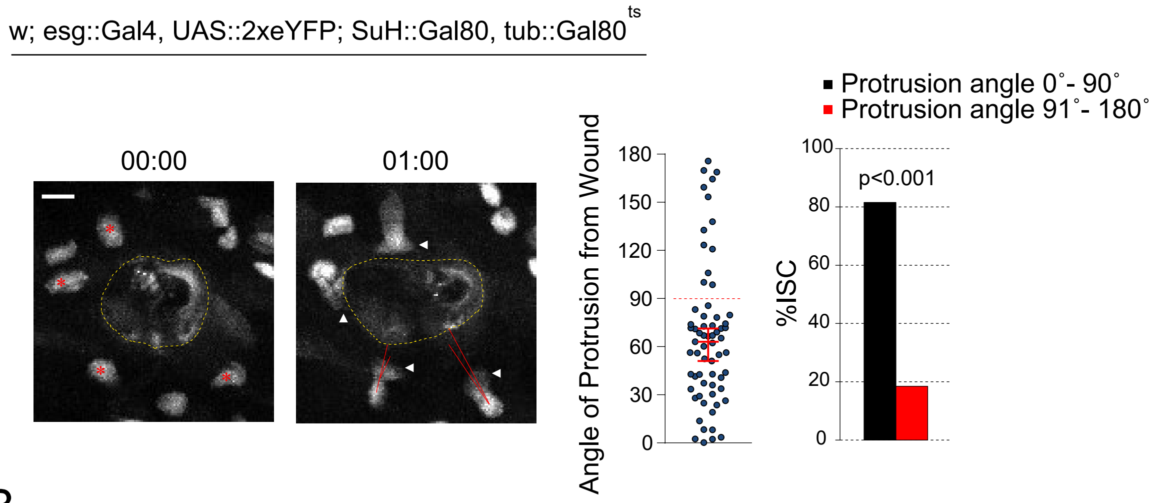


B

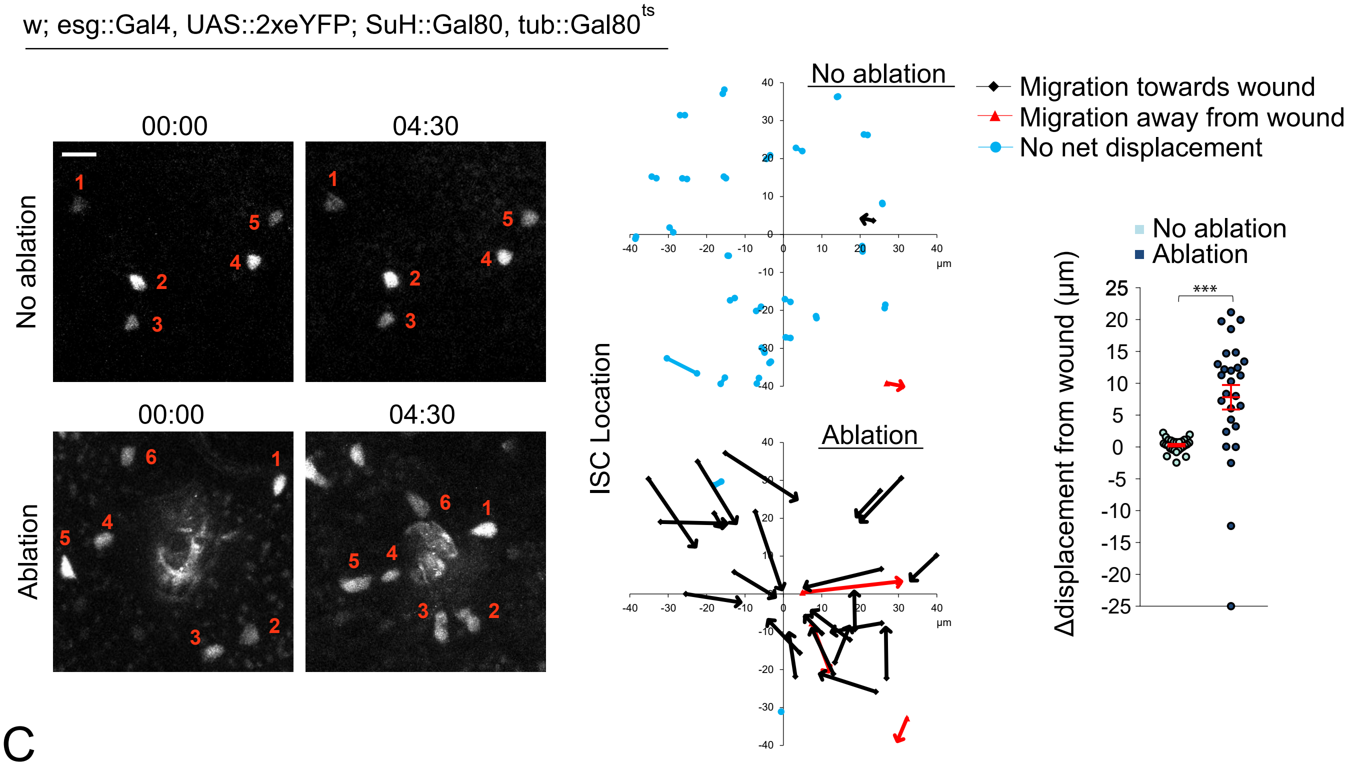


C

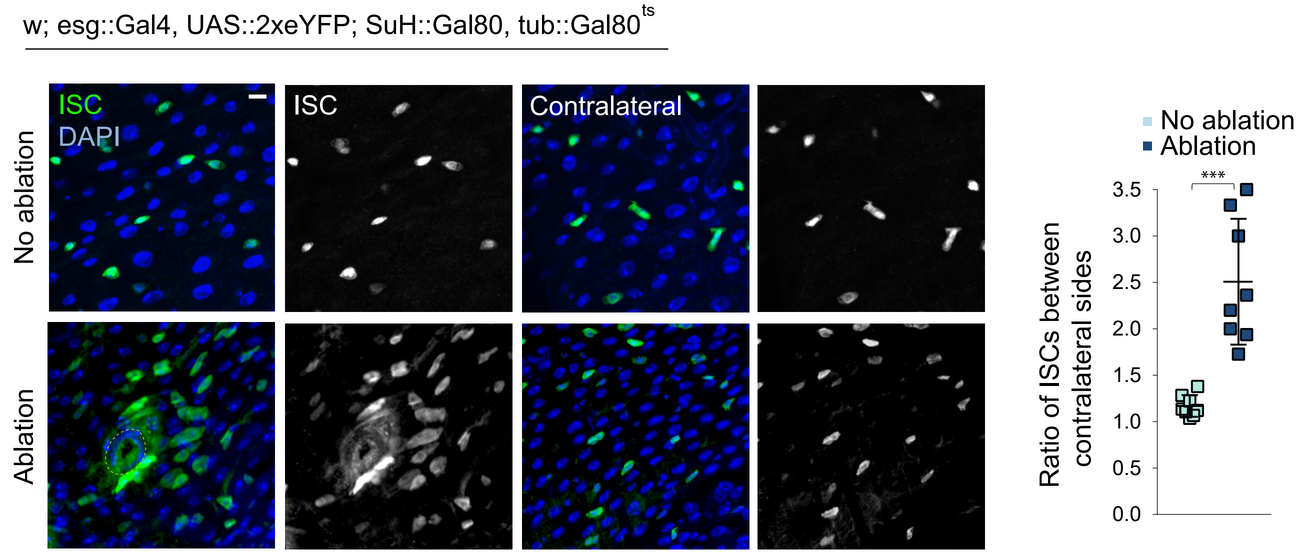

**Supplemental Figure 2: Directionality of ISC migration towards sites of injury, Related to Figure 1.**

A. Montage and quantification of ISC protrusion angle in relation to the wound in ablated tissue. Red asterisk indicate ISCs that will form protrusions after 1hr. Red line depict angle of protrusion in relation to wound, with 0° indicating a protrusion forming along the shortest distance towards the wound. Angles are measured between the vector of the cell body to the middle of the protrusion with the vector of the cell body to the closest point of the wound periphery. B. Montage and quantification of ISC position and change of displacement from the wound immediately following laser ablation (start point) and 4.5hrs later (end point). Numbers in montage correlate to the corresponding ISC at start and end points. The origin of the arrow in the Cartesian plane indicates the ISC start point and the tip of the arrow indicates the ISC end point. x,y coordinates of ISC position in unablated tissue is normalized to an arbitrary point in the middle of the gut, which is set at 0,0. x,y coordinates of ISC position in ablated tissue is normalized to the closest point to the wound periphery of a given ISC, which is set at 0,0. Delta displacement is the difference in distance between the start point and the Cartesian origin, and the end point and the Cartesian origin. C. ISCs accumulated at the site of injury at 4.5hrs post ablation, with increased ISC number around the wound compared to the same area on the contralateral side. median ± 95% CI (A), mean ± SEM (B), or mean ± SD (C); n=65 cells from 6 flies (A), n=27 cells from ≥4 flies (B), n=8 flies (C); ***$P<0.001$, based on chi-square test (A) and one-way ANOVA with Tukey test (B,C). Scale bar = 10µm. Timestamp indicated as hours:minutes.

Supplemental Figure 3

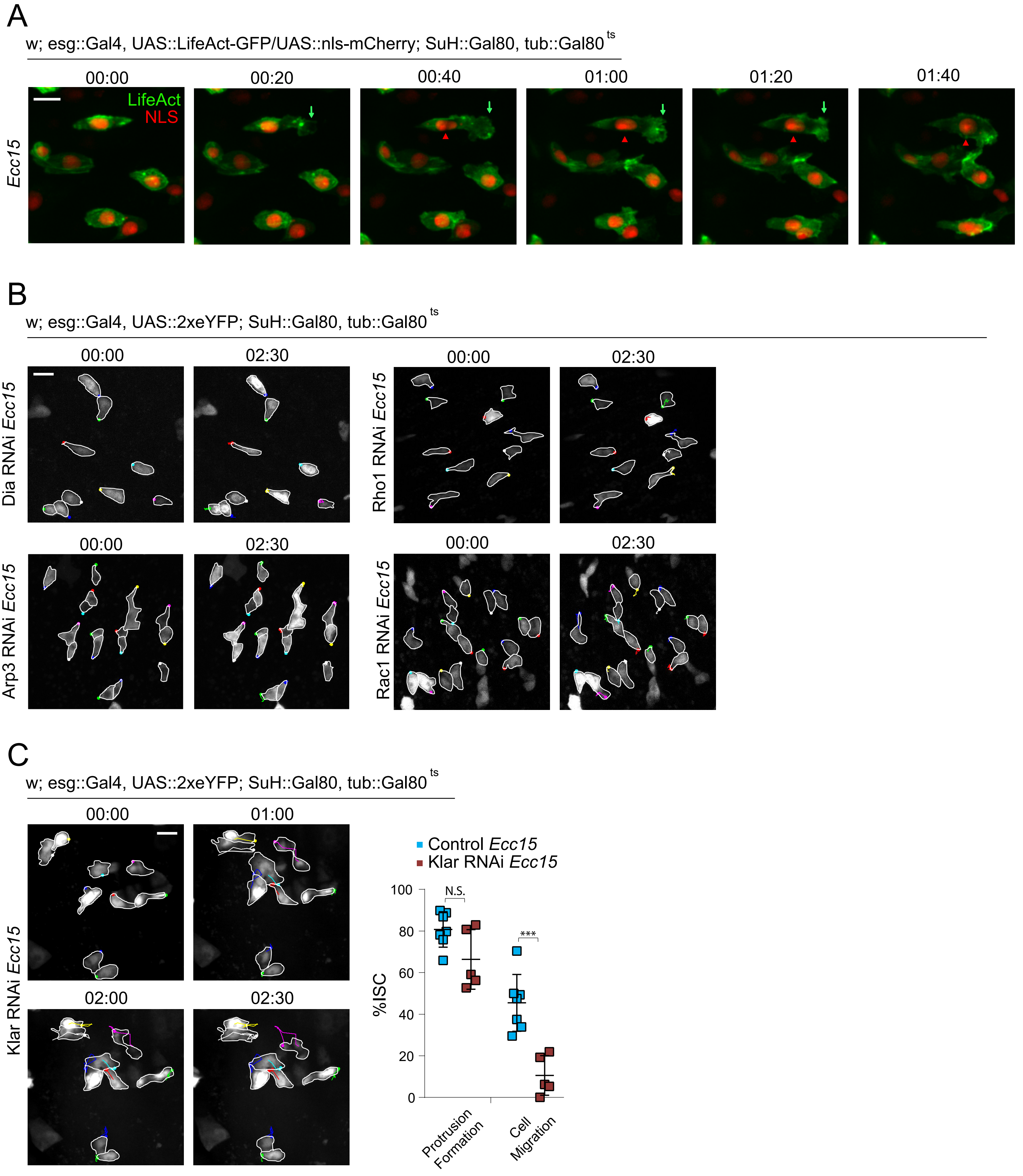

**Supplemental Figure 3: Additional analyses of cytoskeleton-associated proteins during ISC migration, Related to Figure 1.**

A. Montage of ISC migration in *Ecc15*-infected intestines. Green arrows depict leading edge, and red arrowheads depict migrating nucleus. B. Montage of ISC migratory behavior after disrupting actin assembly in *Ecc15*-infected intestines. ISCs outlined in white, and montage tracks measured from the most distal region of the cell cortex. C. Montage and quantification of ISC migration in *Ecc15*-infected Klar$^{RNAi}$ intestines. ISCs outlined in white, and montage tracks measured from the most distal region of the cell cortex. mean ± SD; n≥5 flies; N.S. = not significant, ***P<0.001, based on Student's t-test. Scale bar = 10µm. Timestamp indicated as hours:minutes.

### Supplemental Figure 4

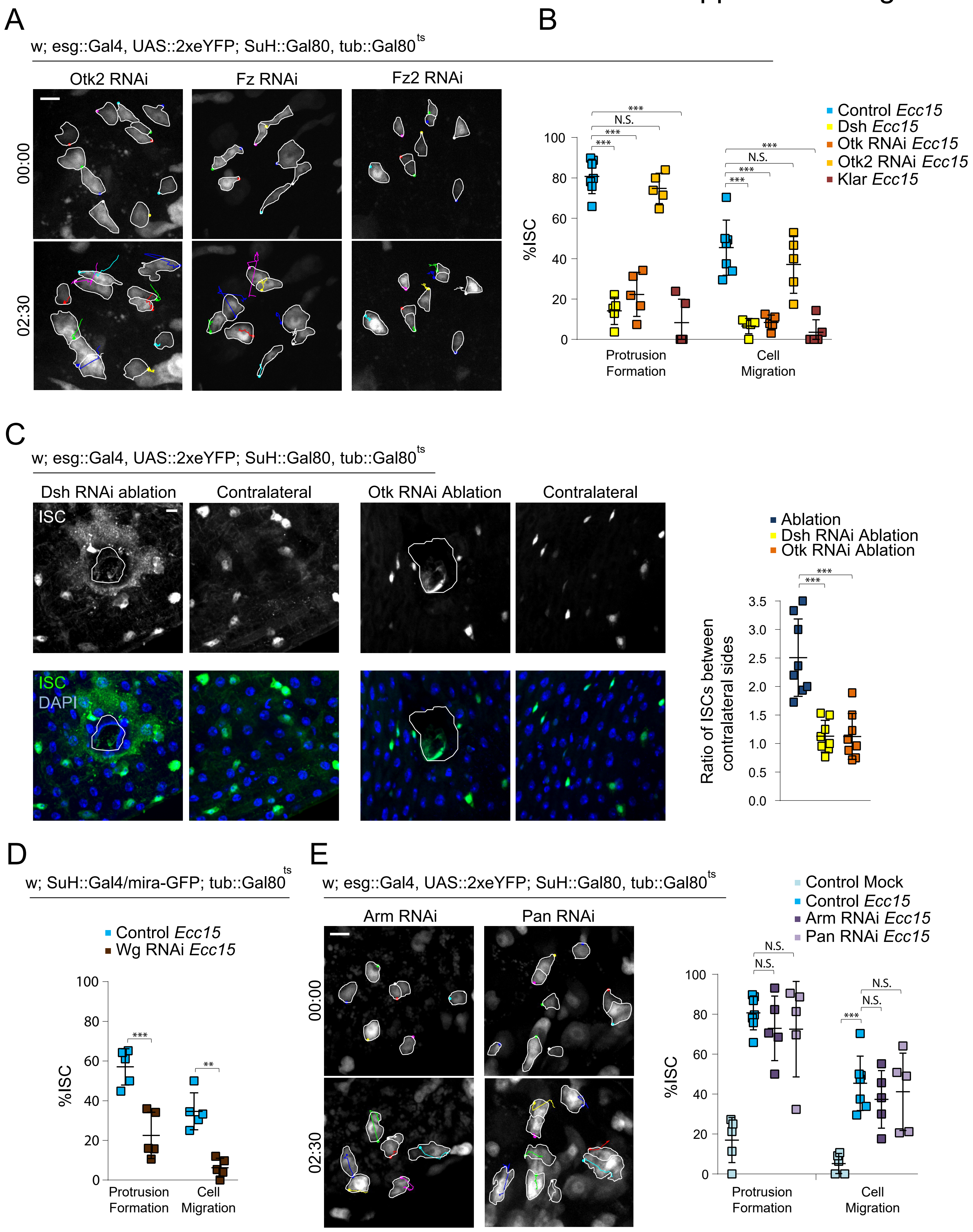

**Supplemental Figure 4: Additional analyses of Wnt signaling roles during ISC migration, Related to Figure 2.**

A. Montage of ISC migration after genetic perturbation in *Ecc15*-infected intestines. ISCs outlined in white, and montage tracks measured from the leading edge or, in cells that did not form protrusions, the most distal region of the cell cortex. B. In *Ecc15*-infected intestines, expression of alternate RNAi against Dsh, Klar, and Otk impaired ISC migration while alternate RNAi against Otk2 had no effect. *Ecc15*-infected controls were taken from Figure 1B, as they served as genetic controls. C. ISCs from $Dsh^{RNAi}$ or $Otk^{RNAi}$ intestines no longer accumulated at the periphery of the wound (outline) 4.5hrs post ablation. D. Depleting Wg specifically in EBs impaired ISC migration. E. Montage and quantification of ISC migratory behavior in *Ecc15*-infected flies with disrupted canonical Wnt signaling. ISCs outlined in white, and montage tracks measured from the leading edge or, in cells that did not form protrusions, the most distal region of the cell cortex. mean ± SD; n≥5 flies (B,D,E), n≥8 flies (C); N.S. = not significant, $^{**}P<0.01$, $^{***}P<0.001$, based on one-way ANOVA with Tukey test (B,C,E) and Student's t-test (D). Scale bar = 10µm. Timestamp indicated as hours:minutes.

### Supplemental Figure 5

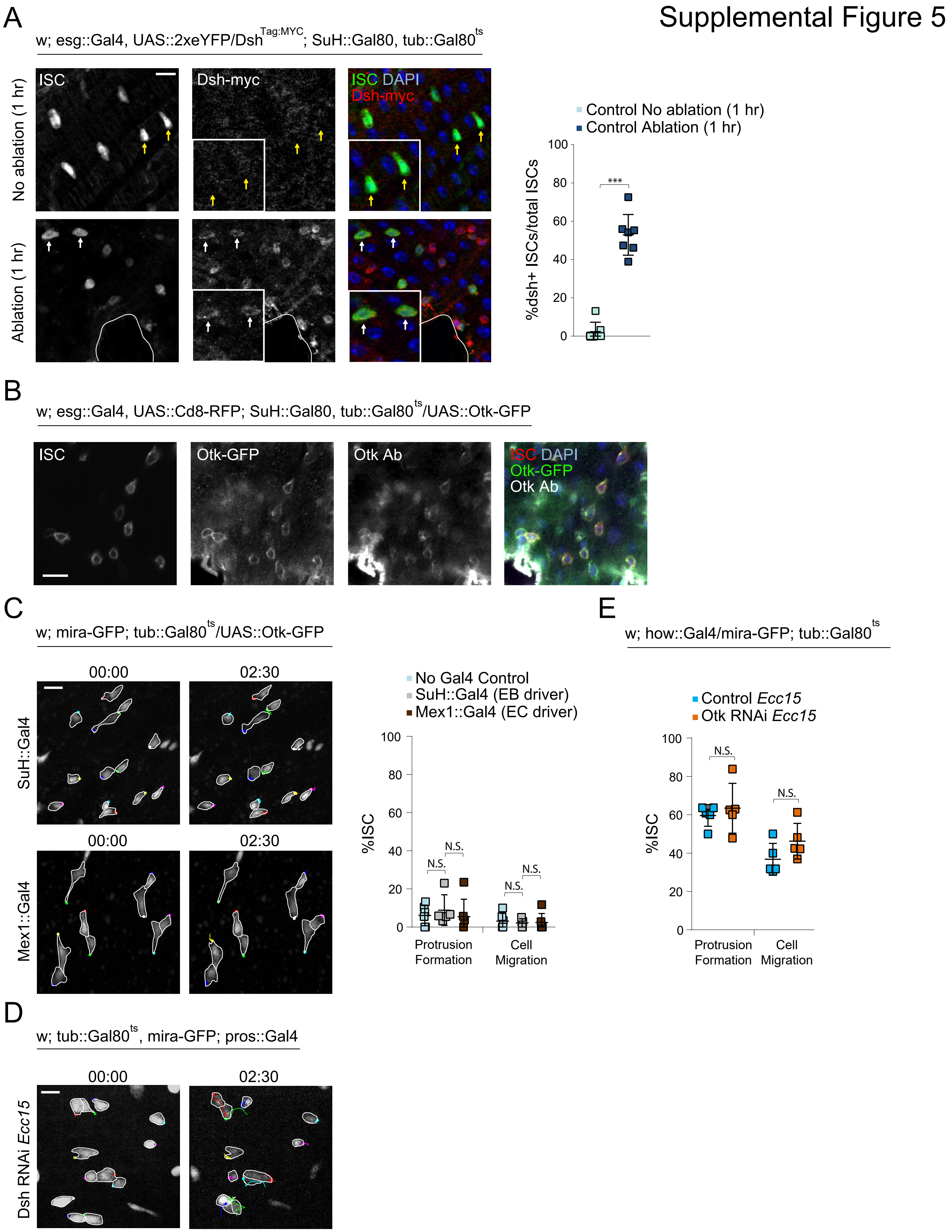

**Supplemental Figure 5: Controls for Dsh and Otk localization, and the effect of additional genetic perturbations in non-ISCs on migration, Related to Figure 3.**

A. Staining for endogenously myc-tagged Dsh reveals cortical decoration in ISCs after 1hr post ablation. White arrows indicate Dsh+ ISCs and yellow arrows indicate Dsh- ISCs. B. Otk antibody co-localizes with GFP-tagged Otk expression. C. Montage and quantification of ISC migration after overexpressing Otk in EBs and ECs. ISCs outlined in white, and montage tracks measured from the most distal region of the cell cortex. D. Montage of ISC migration after depleting Dsh in EEs. ISCs outlined in white, and montage tracks measured from the leading edge or, in cells that did not form protrusions, the most distal region of the cell cortex. E. Depleting Otk in the muscle layer did not significantly alter ISC migration. mean ± SD; n≥5 flies; N.S. = not significant, ***$P<0.001$, based on Student's t-test (A,E) and one-way ANOVA with Tukey test (C). Scale bar = 10µm. Timestamp indicated as hours:minutes.

### Supplemental Figure 6

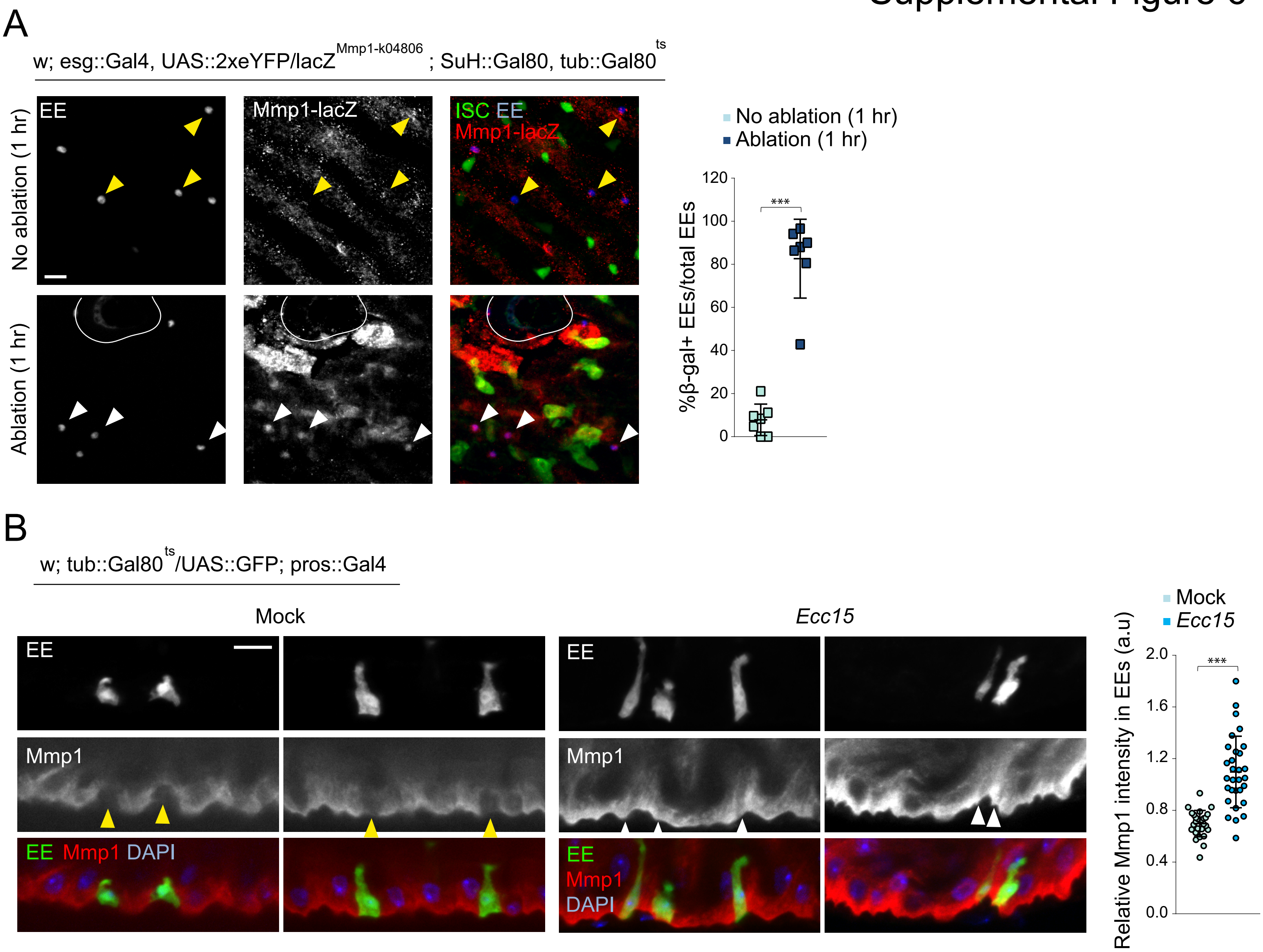

**Supplemental Figure 6: Additional analyses of Mmp1 activity after tissue damage, Related to Figure 4.**

A. Staining for beta Galactosidase in Mmp1-lacZ reporter lines reveal Mmp1 expression in EEs after 1hr post laser ablation. White arrowheads indicate β-gal+ EEs and yellow arrowheads indicate β-gal- EEs. White outline indicates ablation site. B. Mmp1 levels, as determined by immunostaining, increased in EEs after tissue damage. White arrowheads indicate EEs with high Mmp1 levels and yellow arrowheads indicate EEs with low Mmp1 levels. mean ± SD; n=7 flies (A), n≥27 cells from ≥6 flies (B); ***P<0.001, based on Student's t-test. Scale bar = 10μm.

Supplemental Figure 7

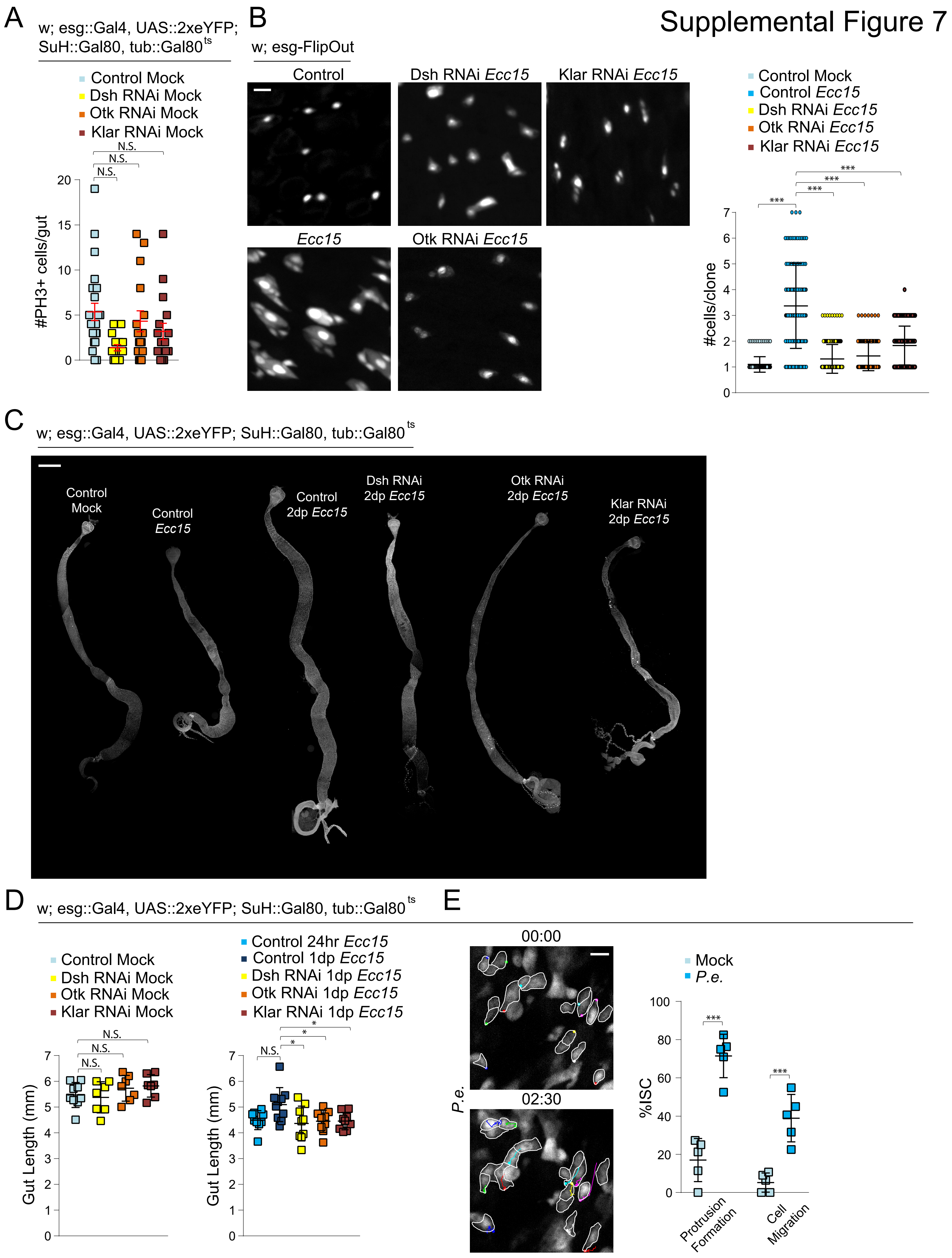

**Supplemental Figure 7: Additional effects of impairing ISC migration on tissue physiology, and migratory behavior after *P.e.* infection, Related to Figure 6.**

A. Quantification of mitotic numbers after disrupting ISC migration in intestines under homeostatic conditions. B. esg-FlipOut clones increased in size after 24hr *Ecc15* infection. This increase in clone size was stunted when ISC migration was impaired. C. Representative images of full-length intestines. D. Gut length after impairing ISC migration in homeostatic condition and one day of recovery after 24hr *Ecc15* infection. E. Montage and quantification of migratory behavior after *P.e.* infection. ISCs outlined in white, and montage tracks measured from the leading edge or, in cells that did not form protrusions, the most distal region of the cell cortex. Mock-treated controls were taken from Figure 1B. mean ± SEM (A) or mean ± SD (B,D,E); n≥15 flies (A), n≥129 clones from 5 flies (B), n≥7 flies (D), n=5 flies (E); N.S. = not significant, *P<0.05, ***P<0.001, based on one-way ANOVA with Tukey test (A,B,D) and Student's t-test (E). Scale bar = 10µm (B,E), 500µm (C). Timestamp indicated as hours:minutes.

**Supplemental Video Legends**

**Supplemental Video 1: ISC migratory behavior during homeostasis, Related to Figure 1.**

Cytoplasmic GFP and mCherry-tagged NLS was expressed in ISCs for four days. The fly was fed 5% sucrose for 16 hours, before the intestine dissected and imaged *ex vivo* at 10-min intervals. Time stamp indicated as hours:minutes:seconds. Video dimensions are 63µm x 63µm (width x height). Tracks measured from the center of the nucleus.

**Supplemental Video 2: ISC migratory behavior after *Ecc15* infection, Related to Figure 1.**

Cytoplasmic GFP and mCherry-tagged NLS was expressed in ISCs for four days. The fly was infected with *Ecc15* for 16 hours, before the intestine were dissected and imaged *ex vivo* at 10-min intervals. Time stamp indicated as hours:minutes:seconds. Video dimensions are 63µm x 63µm (width x height). Tracks measured from the center of the nucleus.

**Supplemental Video 3: EB migratory behavior after *Ecc15* infection, Related to Figure 1.**

Cytoplasmic GFP was expressed in EBs for four days. The fly was infected with *Ecc15* for 16 hours, before the intestine was dissected and imaged *ex vivo* at 10-min intervals. Tracks were measured at the most distal region of the cell cortex. Time stamp indicated as hours:minutes:seconds. Video dimensions are 81µm x 81µm (width x height).

**Supplemental Video 4: EE migratory behavior after *Ecc15* infection, Related to Figure 1.**

Cytoplasmic GFP was expressed in EEs for four days. The fly was infected with *Ecc15* for 16 hours, before the intestine was dissected and imaged *ex vivo* at 10-min intervals. Tracks were measured at the most distal region of the cell cortex. Time stamp indicated as hours:minutes:seconds. Video dimensions are 81µm x 81µm (width x height).

**Supplemental Video 5: ISC migratory behavior in undamaged intestine, Related to Figure 1.**

Cytoplasmic GFP was expressed in ISCs for four days. Undamaged intestine was imaged *ex vivo* at 10-min intervals on the 2-photon microscopy system, and served as a control for ablated intestines. Minimal protrusion formations occurred. Time stamp indicated as hours:minutes:seconds. Video dimensions are 137µm x 137µm (width x height).

**Supplemental Video 6: ISC migratory behavior after ablation, Related to Figure 1.**

Cytoplasmic GFP was expressed in ISCs for four days. Intestine was dissected and imaged *ex vivo* at 10-min intervals. Laser ablation occurred 30" before the second time point, resulting in multiple ISCs exhibiting protrusion formation throughout the remainder of the capture. Time stamp indicated as hours:minutes:seconds. Video dimensions are 137µm x 137µm (width x height).

**Supplemental Video 7: The actin cytoskeleton is largely static under homeostatic conditions, Related to Figure 1.**

LifeAct-GFP and mCherry-tagged NLS was expressed in ISCs for four days. The fly was fed 5% sucrose for 16 hours, before the intestine was dissected, treated with 0.1% DMSO for 30', and imaged ex vivo at 10-min intervals. Time stamp indicated as hours:minutes:seconds. Video dimensions are 63µm x 63µm (width x height).

**Supplemental Video 8: Actin dynamics greatly increase after *Ecc15* infection, Related to Figure 1.**

LifeAct-GFP and mCherry-tagged NLS was expressed in ISCs for four days. The fly was infected with *Ecc15* for 16 hours, before the intestine was dissected, treated with 0.1% DMSO for 30', and imaged *ex vivo* at 10-min intervals. Time stamp indicated as hours:minutes:seconds. Video dimensions are 63µm x 63µm (width x height).

**Supplemental Video 9: Treatment with Cytochalasin B abolishes actin dynamics in *Ecc15*-infected intestine, Related to Figure 1.**

LifeAct-GFP was expressed in ISCs for four days. The fly was infected with *Ecc15* for 16 hours, before the intestine was dissected, treated with 10μM Cytochalasin B for 30', and imaged *ex vivo* at 10-min intervals. Time stamp indicated as hours:minutes:seconds. Video dimensions are 33μm x 39μm (width x height).

**Supplemental Video 10: Arp3 RNAi impairs ISC migratory behavior in *Ecc15*-infected intestine, Related to Figure 1.**

Arp3 RNAi and cytoplasmic GFP was expressed in ISCs for four days. The fly was infected with *Ecc15* for 16 hours, before the intestine was dissected and imaged *ex vivo* at 10-min intervals. Tracks were measured from the edge of the protrusion, or, for ISCs that did not exhibit protrusion formation, the most distal region of the cell cortex. Time stamp indicated as hours:minutes:seconds. Video dimensions are 81μm x 81μm (width x height).

**Supplemental Video 11: Klar RNAi impairs cell body translocation but not protrusion formation in *Ecc15*-infected intestine, Related to Figure 1.**

Klar RNAi and cytoplasmic GFP was expressed in ISCs for four days. The fly was infected with *Ecc15* for 16 hours, before the intestine was dissected and imaged *ex vivo* at 10-min intervals. Tracks were measured from the edge of the protrusion, or, for ISCs that did not exhibit protrusion formation, the most distal region of the cell cortex. Time stamp indicated as hours:minutes:seconds. Video dimensions are 81μm x 81μm (width x height).

**Supplemental Video 12: Dsh RNAi impairs ISC migratory behavior in *Ecc15*-infected intestine, Related to Figure 2.**

Dsh RNAi and cytoplasmic GFP was expressed in ISCs for four days. The fly was infected with *Ecc15* for 16 hours, before the intestine was dissected and imaged *ex vivo* at 10-min intervals. Tracks were measured from the edge of the protrusion, or, for ISCs that did not exhibit protrusion formation, the most distal region of the cell cortex. Time stamp indicated as hours:minutes:seconds. Video dimensions are 81μm x 81μm (width x height).

**Supplemental Video 13: Otk RNAi impairs ISC migratory behavior in *Ecc15*-infected intestine, Related to Figure 2.**

Otk RNAi and cytoplasmic GFP was expressed in ISCs for four days. The fly was infected with *Ecc15* for 16 hours, before the intestine was dissected and imaged *ex vivo* at 10-min intervals. Tracks were measured from the edge of the protrusion, or, for ISCs that did not exhibit protrusion formation, the most distal region of the cell cortex. Time stamp indicated as hours:minutes:seconds. Video dimensions are 81µm x 81µm (width x height).

**Supplemental Video 14: Dsh RNAi impairs ISC migratory behavior in ablated intestine, Related to Figure 2.**

Dsh RNAi and cytoplasmic GFP was expressed in ISCs for four days. Intestine was dissected and imaged *ex vivo* at 10-min intervals. Laser ablation occurred 30" before the second time point. Time stamp indicated as hours:minutes:seconds. Video dimensions are 137µm x 137µm (width x height).

**Supplemental Video 15: Otk RNAi impairs ISC migratory behavior in *Ecc15*-infected intestine, Related to Figure 2.**

Otk RNAi and cytoplasmic GFP was expressed in ISCs for four days. Intestine was dissected and imaged *ex vivo* at 10-min intervals. Laser ablation occurred 30" before the second time point. Time stamp indicated as hours:minutes:seconds. Video dimensions are 137µm x 137µm (width x height).

**Supplemental Video 16: Treatment with LGK974 abolishes ISC migratory behavior in ablated intestine, Related to Figure 2.**

Cytoplasmic GFP was expressed in ISCs for four days. Intestine was dissected, treated with 1µM LGK974 B for 30', and imaged *ex vivo* at 10-min intervals. Laser ablation occurred 30" before the second time point. Time stamp indicated as hours:minutes:seconds. Video dimensions are 137µm x 137µm (width x height).

**Supplemental Video 17: Otk OE promotes protrusion dynamics, Related to Figure 2.**

Otk-GFP and cytoplasmic GFP was expressed in ISCs for four days. Intestine was dissected and imaged *ex vivo* at 10-min intervals. Time stamp indicated as hours:minutes:seconds. Video dimensions are 39µm x 33µm (width x height).

**Supplemental Video 18: Otk OE in EEs promotes migratory behavior in ISCs, Related to Figure 3.**

Otk was overexpressed in EEs for four days, and ISCs were specifically labelled with mira-GFP. Intestine was dissected and imaged *ex vivo* at 10-min intervals. Tracks were measured from the edge of the protrusion, or, for ISCs that did not exhibit protrusion formation, the most distal region of the cell cortex. Time stamp indicated as hours:minutes:seconds. Video dimensions are 81µm x 81µm (width x height).

**Supplemental Video 19: Otk RNAi in EEs impairs migratory behavior in ISCs in *Ecc15*-infected intestine, Related to Figure 3.**

Otk RNAi was expressed in EEs for four days, and ISCs were specifically labelled with mira-GFP. The fly was infected with *Ecc15* for 16 hours, before the intestine was dissected and imaged *ex vivo* at 10-min intervals. Tracks were measured from the edge of the protrusion, or, for ISCs that did not exhibit protrusion formation, the most distal region of the cell cortex. Time stamp indicated as hours:minutes:seconds. Video dimensions are 81µm x 81µm (width x height).

**Supplemental Video 20: Treatment with Ptk7 promotes ISC migratory, Related to Figure 4.**

Lifeact-GFP was expressed in ISCs for four days. Intestine was dissected, treated with 1µg/ml N-terminal Ptk7 for 30’, and imaged *ex vivo* at 10-min intervals. Time stamp indicated as hours:minutes:seconds. Video dimensions are 39µm x 33µm (width x height).

**Supplemental Video 21: Otk RNAi abolishes protrusion formation in Ptk7-treated intestine, Related to Figure 4.**

Otk RNAi and Lifeact-GFP was expressed in ISCs for four days. Intestine was dissected, treated with 1µg/ml N-terminal Ptk7 for 30', and imaged *ex vivo* at 10-min intervals. Time stamp indicated as hours:minutes:seconds. Video dimensions are 39µm x 33µm (width x height).

**Supplemental Video 22: Timp OE in EEs impairs ISC migratory behavior in *Ecc15*-infected intestine, Related to Figure 4.**

Timp was overexpressed in EEs for four days, and ISCs were specifically labelled with mira-GFP. The fly was infected with *Ecc15* for 16 hours, before the intestine was dissected and imaged *ex vivo* at 10-min intervals. Time stamp indicated as hours:minutes:seconds. Video dimensions are 81µm x 81µm (width x height).

**Supplemental Video 23: Treatment with TAPI-1 abolishes ISC migratory behavior in ablated intestine, Related to Figure 4.**

Cytoplasmic GFP was expressed in ISCs for four days. Intestine was dissected, treated with 20µM TAPI-1 for 30', and imaged *ex vivo* at 10-min intervals. Laser ablation occurred 30" before the second time point. Time stamp indicated as hours:minutes:seconds. Video dimensions are 137µm x 137µm (width x height).

**Supplemental Video 24: Wee1 OE impairs ISC migratory behavior in *Ecc15*-infected intestine, Related to Figure 6.**

Wee1 and cytoplasmic GFP was overexpressed in ISCs for four days. The fly was infected with *Ecc15* for 16 hours, before the intestine was dissected and imaged *ex vivo* at 10-min intervals. Tracks were measured from the edge of the protrusion, or, for ISCs that did not exhibit protrusion formation, the most distal region of the cell cortex. Time stamp indicated as hours:minutes:seconds. Video dimensions are 81µm x 81µm (width x height).

**Supplemental Video 25: Wee1 RNAi promotes ISC migratory behavior, Related to Figure 6.**

Wee1 RNAi and cytoplasmic GFP was expressed in ISCs for four days. The intestine was dissected and imaged *ex vivo* at 10-min intervals. Tracks were measured from the edge of the protrusion, or, for ISCs that did not exhibit protrusion formation, the most distal region of the cell cortex. Time stamp indicated as hours:minutes:seconds. Video dimensions are 81µm x 81µm (width x height).
